# Modular functional brain network organization contributes to training-related changes in task switching in children

**DOI:** 10.64898/2026.02.26.708262

**Authors:** Sina A. Schwarze, Ulman Lindenberger, Silvia A. Bunge, Yana Fandakova

## Abstract

Cognitive training often aims to improve cognitive skills, but improvements in cognitive performance have been variable. One factor that has been found to predict training outcomes is the degree of modularity of functional brain networks, defined as the extent to which brain regions are more strongly connected to regions within the same functional subnetwork than to regions outside of the subnetwork. Specifically, more modular organization of functional brain networks at baseline has been associated with greater benefits from cognitive training in adults. During childhood, cognitive development is marked by a slow progression towards network integration and segregation, which together contribute to increasing modularity. Thus, network modularity might also be an important predictor of training outcomes in children. To investigate whether individual differences in network modularity predict outcomes of task-switching training in children, we examined 87 children aged 8 to 11 years who completed nine weeks of varying amounts of task-switching training. Prior to training, children showed lower network modularity than adults, in line with previously reported developmental changes in network configuration. More modular organization of functional brain networks before training was associated with higher overall accuracy during task switching and with faster improvements in integrated speed-accuracy scores, especially at the beginning of training. These results suggest that more modular functional networks might allow for faster adaptation to training demands in children and thus faster improvements with training.

**Graphical abstract:** 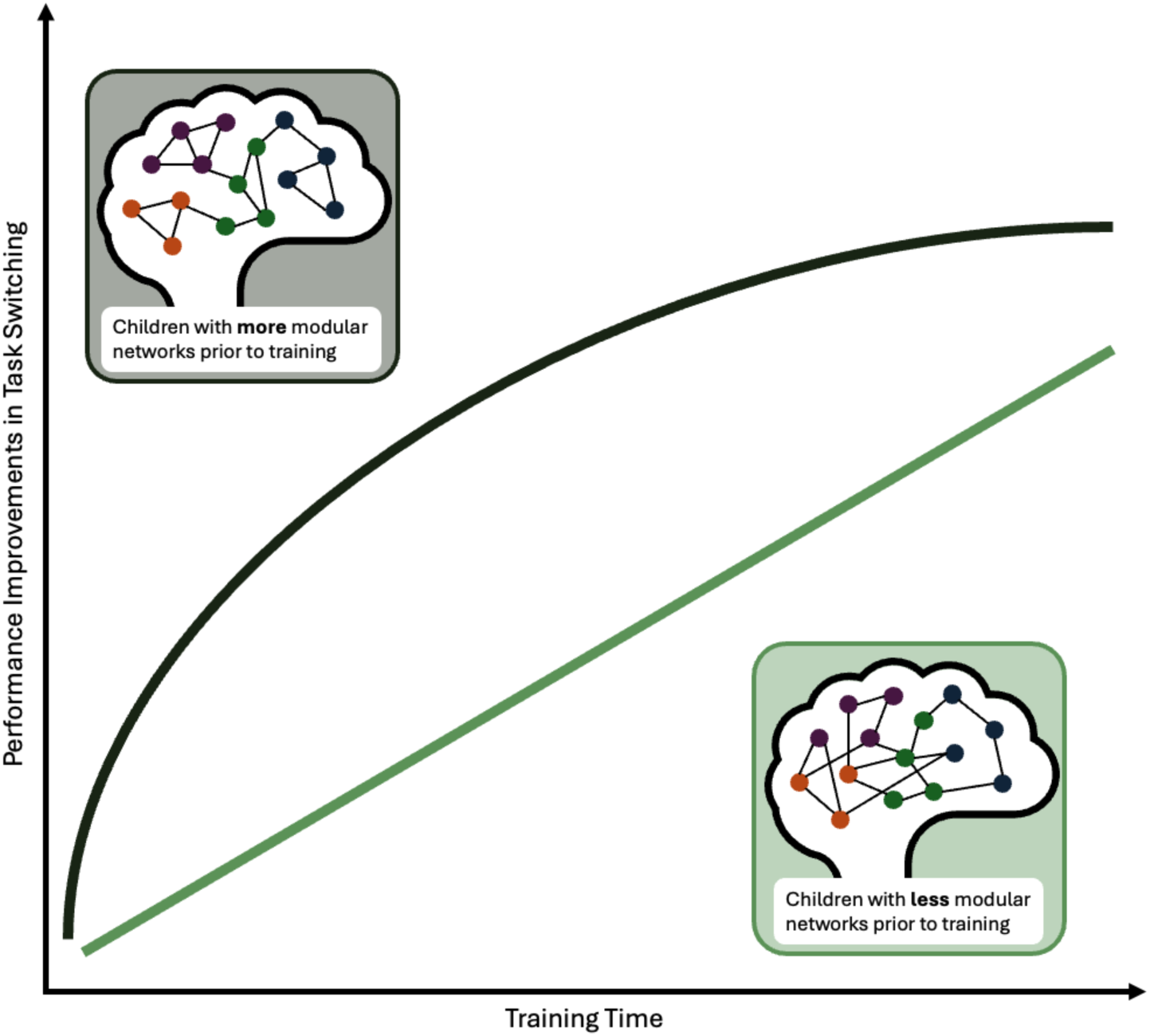

**Highlights:**

- Children showed less modular network organization than adults.
- More modular networks before training were associated with overall higher accuracy.
- Children with more modular networks adapted more quickly to training demands.

## 1. Introduction

Executive functions, a set of processes coordinating cognition and behavior in the pursuit of goals, continue to develop throughout childhood and adolescence (Tervo-Clemmens et al., 2023) in association with positive real-life outcomes (Best et al., 2011; Moffitt et al., 2011; Titz and Karbach, 2014). Accordingly, there has been increasing interest in identifying training programs that improve executive functions in children (Kray and Dörrenbächer, 2020; Rueda et al., 2005; Zhang et al., 2019). However, training studies have shown varying degrees of success in improving executive functioning, with large individual differences in training outcomes among individuals undergoing the same training protocol (Jolles and Crone, 2012; Katz et al., 2021; Kray and Dörrenbächer, 2020). It is therefore important to examine factors that can help predict who will benefit most from a particular training program (Brehmer et al., 2007; Fandakova et al., 2012; Karbach et al., 2017; Reuter-Lorenz and Lustig, 2005). In the present study we explore how characteristics of functional brain network organization predict individual differences in the outcome of task-switching training in children.

### 1.1 Development of task switching and underlying brain networks

Generally, children show lower performance on executive functions tasks than adults (Diamond, 2013; Luna, 2009; Tervo-Clemmens et al., 2023). One core aspect of executive functions is shifting or switching, denoting the ability to adapt to changes in a task’s rules or goals (Diamond, 2013; Miyake et al., 2000). Shifting is often measured using task-switching paradigms that require participants to switch between different tasks in intermixed task blocks (Monsell, 2003). Task-switching paradigms often also include single-task blocks that require participants to perform each task separately (see Koch and Kiesel, 2022, for a review).

The demands of switching between different tasks in the task-switching paradigm generally result in two types of performance costs. First, mixing costs are elicited by greater demands on maintaining, managing, and monitoring multiple task rules in blocks of intermixed tasks as opposed to single-task blocks (e.g., Allport et al., 1994; Jersild, 1927; Rubin and Meiran, 2005). Second, switch costs result from a greater demand to retrieve and update the new task rule and to inhibit the no-longer relevant task rule on switch compared to repeat trials in intermixed blocks (e.g., Allport et al., 1994; Mayr and Kliegl, 2000; Meiran, 1996; Rogers and Monsell, 1995; Wylie and Allport, 2000). Notably, while both mixing and switch costs decrease throughout childhood and adolescence, children show more prolonged development of managing and monitoring multiple task rules as evident in larger age differences in mixing costs that last well into adolescence (Cragg and Chevalier, 2012; Gupta et al., 2009; Huizinga et al., 2006; Huizinga and van der Molen, 2007; Schwarze et al., 2024, 2023). Given the protracted development of task switching and its relevance for outcomes such as academic achievement and well-being (Liew, 2012; Titz and Karbach, 2014; Yeniad et al., 2013), there is increased interest in understanding the mechanisms underlying individual differences in task-switching development. To this end, examining individual differences in the trajectories and outcomes of task-switching training offers a unique window into the plasticity of the developing brain.

Developmental improvements in executive functions, and task switching in particular, have been linked to various aspects of brain development, including structural development of the prefrontal cortex (Sydnor et al., 2021), a more refined recruitment of frontoparietal brain regions (Chevalier et al., 2019; Church et al., 2017; Crone et al., 2006; Schwarze et al., 2023; Wendelken et al., 2012), and the development of large-scale brain networks (Bassett et al., 2018; Cui et al., 2020; Grayson and Fair, 2017; Gu et al., 2020; Kupis et al., 2021; Marek et al., 2015; Pines et al., 2022; Tooley et al., 2022b). Additionally, individual differences in cognitive flexibility among children have been linked to variation in how functional networks interact during task performance (Kupis et al., 2021; Kupis and Uddin, 2023), stressing the relevance of network organization for the development of children’s task-switching ability.

Generally, child development is marked by protracted network integration and segregation, as measured via magnetic resonance imaging (MRI; Baum et al., 2017; Fair et al., 2007; Keller et al., 2023; Luna et al., 2015; Sun et al., 2024; Yin et al., 2025), reflecting the ongoing specialization of brain networks over the course of development. Integration refers to strengthening of connectivity among regions of the same subnetwork, whereas segregation refers to weakening of connectivity among regions of different subnetworks.

A graph theoretic metric referred to as modularity can be used to quantify the extent of integration and segregation of a network, such that a network is described as more modular, the more integrated and segregated it is (Bullmore and Bassett, 2011; Newman and Girvan, 2004; Sporns and Betzel, 2016). In a modular network, subnetworks are strongly connected within themselves and clearly distinguishable from other subnetworks. In adults, more modular networks have been associated with better learning (Bassett et al., 2015; Ellefsen et al., 2015) and better working memory (Braun et al., 2015; Stevens et al., 2012). Over the course of child and adolescent development, the functional networks commonly identified in adults become increasingly modular with age (McCormick et al., 2021; Wang et al., 2020) and are closely associated with improvements in executive functions (Baum et al., 2017; Wang et al., 2020).

### 1.2 Task-switching training and individual differences in training outcomes

Studies investigating the effects of task-switching training typically compare a group that practices switching between tasks (i.e., task-switching training) to a group that performs the same tasks as single-task blocks for the same training duration. These studies have demonstrated that task-switching training can lead to improved performance and lower mixing and switch costs, especially on the trained task-switching paradigms and on untrained paradigms that closely resemble the trained tasks and that target task-switching abilities in a similar manner (Berryhill and Hughes, 2009; Cepeda et al., 2001; Dörrenbächer et al., 2014; Kray and Lindenberger, 2000; Minear and Shah, 2008; Strobach et al., 2012; von Bastian and Oberauer, 2013). While an exhaustive review on task-switching training studies across the lifespan is beyond the scope of the present paper, we refer to Kray and Dörrenbächer (2020) for such a detailed review.

However, while training effects have been documented based on comparisons between training groups, it has also become evident that individuals differ substantially in their benefit from task-switching and, likewise, cognitive training in other domains (Buttelmann and Karbach, 2017; Feng et al., 2023; Karbach et al., 2017; Katz et al., 2021; Könen and Karbach, 2015). For instance, previous studies have consistently demonstrated that participants’ baseline performance (i.e., prior to training) is associated with training outcomes (Borella et al., 2017; Karbach et al., 2017; Katz et al., 2021; Lövdén et al., 2012).

However, this pattern is not uniform across training studies: in some studies individuals with better baseline performance benefit more from training (Foster et al., 2017; Rhodes and Katz, 2017), while in other studies individuals with poorer performance benefit more (Carretti et al., 2017; Karbach et al., 2017). Matching the latter pattern, studies investigating age differences in task-switching training suggest that training-related improvements in childhood are more prominent in mixing costs potentially due to their ongoing development offering more room for improvement (Karbach and Kray, 2009; Schwarze et al., 2025). These studies have also revealed that children benefit more from training than adults, especially on untrained task-switching paradigms (Cepeda et al., 2001; Karbach et al., 2017; Karbach and Kray, 2009). However, the mechanisms that contribute to individual differences in outcomes of task-switching training in children are still largely unclear (Buttelmann and Karbach, 2017).

The observation of substantial individual differences in training outcomes is as old as cognitive training research itself (Dallenbach, 1919, 1914), and designing effective training programs requires a closer consideration of the complexity of factors contributing to individual differences (Katz et al., 2021; Ørskov et al., 2021; Smid et al., 2020). Here, we explore the degree to which individual differences in brain network organization may represent a factor contributing to individual differences in task-switching training benefits among children. To this end, we utilize neuroimaging and network modularity as a graph theoretical measure of brain network organization. A brain network approach to individual differences in cognitive training might be particularly relevant, given that learning, as a process of integrating different types of information, involves many brain regions and their interactions (Bassett et al., 2011; Gerraty et al., 2018, 2014; McCormick et al., 2021) and that these interactions during task-switching differ in childhood (Kupis et al., 2021; Kupis and Uddin, 2023).

### 1.3 Network modularity as predictor of training outcomes

Network modularity has been proposed as a measure to predict training outcomes based on network connectivity (Gallen and D’Esposito, 2019). More specifically, it has been theorized that more modular networks are closer to the ideal state of network architecture for task execution and thus enable a more optimal state to approach training demands. Higher modularity may allow for more effective information abstraction that supports faster processing and exhibits greater adaptability to changing environmental demands (Kashtan and Alon, 2005). Accordingly, studies in younger and older adults have demonstrated that individuals showing greater brain network modularity at baseline benefit more from interventions targeting cognitive skills after accounting for differences in baseline ability (Baniqued et al., 2019; Bassett et al., 2011; Ellefsen et al., 2015; Gallen et al., 2016). Additionally, previous research has shown that network modularity changes with cognitive training such that functional networks become more segregated (Bassett et al., 2015).

So far, few studies have investigated how the ongoing development of network organization in childhood and adolescence is related to individual differences in training outcomes. One study that implemented a physical activity intervention demonstrated that 8–9-year-old children with more modular network organization prior to the intervention showed greater increases in executive functions after training (Chaddock-Heyman et al., 2020), providing first evidence for the relevance of network modularity for improving executive functions in children. Another study investigated the association between network modularity and short-term learning from performance feedback in a longitudinal sample of participants aged between 8 and 29 years (McCormick et al., 2021). Results of the latter study showed that more modular network organization was associated with greater improvements in learning over time, but only after adolescence, suggesting an intriguing developmental difference in the role of network organization for learning.

### 1.4 Present study

In the present study, we tested whether individual differences in functional brain network modularity were associated with individual differences in the outcomes of an intensive task-switching and single-task training in 87 children aged between 8 and 11 years. We have previously compared group-level differences in this sample (Schwarze et al., 2025). Our results showed training-related decreases in mixing costs, but not in switch costs in the training groups compared to the control group. Moreover, we leveraged additional laboratory assessments during training, observing rapid improvements of accuracy mixing costs after the start of training in both the single-task and task-switching groups, with group differences only becoming apparent in the later phase of training. Specifically, towards the end of training only the task-switching group maintained the training benefits from the first phase (Schwarze et al., 2025).

While these group comparisons demonstrate training-related improvements in task-switching performance on the group level, the magnitude of this improvement differed substantially across individual children, prompting us to further explore potential factors contributing to these individual differences. Given the protracted development of functional brain network organization, network modularity may be especially relevant as a predictor of training-related behavioral plasticity in children (Chaddock-Heyman et al., 2020) or may not yet affect outcomes of cognitive training, as suggested by previous findings on network modularity and learning (McCormick et al., 2021).

Here, we tested two predictions: first, that functional network modularity prior to training and/or changes in functional network modularity with training would be associated with individual differences in changes in task-switching performance with training; and second, that functional brain network modularity would increase with training. We collapsed across the single-task and the task-switching training groups to increase the power for testing the relationship between modularity and individual differences in improvement, since we did not have a specific hypothesis that this association differed between the two groups, and because the two types of training were quite similar (i.e., the single-task training groups also performed some blocks of intermixed tasks in each training session).

## 2. Methods

Before the start of data analysis, hypotheses and analysis plans were preregistered at https://aspredicted.org/vh6x-q55f.pdf. Training-related changes in accuracy, response times (RTs), and task-based activation and connectivity have previously been reported in Schwarze et al. (2025).

### 2.1 Participants

A total of 87 children between the age of 8 and 11 years (M = 9.94 years, SD = 0.7; 43 girls, 44 boys) who underwent four MRI assessments within a task-switching training study were included in the present analyses. Note that additional children were included in the behavioral analyses in Schwarze et al. (2025) – however, they completed the identical task in a mock scanner and thus had no MRI data available. Furthermore, the initial publication included a passive control group that did not undergo training. As we focus on the association of network modularity with changes in performance with training in the present analyses, we excluded the data of the passive control group. To test whether children in the present sample showed the expected pattern of less modular functional brain networks than adults, we compared their modularity at the pre-training session to a sample of young adults (N = 53; mean age = 24.7, SD = 2.6; 28 women, 25 men) who were only scanned once and who did not undergo any training (see Schwarze et al., 2023).

Children were pseudo-randomly assigned to one of two training groups: either practicing mainly single tasking (SI group; N = 44; 21 girls, 23 boys; mean age = 9.98, SD = 0.69) or task switching (SW group; N = 43; 22 girls, 21 boys; mean age = 9.9, SD = 0.71; see details on training in section 2.2). Participants were screened for MRI suitability, were right-handed, spoke German as their primary language, and had no history of neurological or psychological diseases. Parents and children provided informed written consent. All participants were reimbursed with 10 € per hour spent at the laboratory and received an additional bonus of 40 € for the completion of all MRI sessions and training sessions. Children additionally received a toy as a reward for their performance in the training sessions. The study was conducted in line with the Declaration of Helsinki and approved by the ethics committee of the Freie Universität Berlin.

To ensure high data quality, children were excluded from analysis based on session-specific preregistered performance and movement criteria, as well as a minimum criterion for the number of training sessions completed. Specifically, if a child showed below 50% accuracy in the run of single blocks (run 1, see below for more details on the paradigm) or below 35% accuracy in either of the two runs of mixed blocks (runs 2 and 3) in a given session of the task-switching paradigm, their data of that session were excluded from analyses. Session-specific data were further excluded if any of the fMRI runs of a session exceeded 50% of low-quality volumes, i.e., volumes with framewise displacement (Power et al., 2012) above 0.4 mm (cf. Dosenbach et al. 2017). The final sample included 63 children at session A (SW = 33, SI = 30), 56 at session B (SW = 33, SI = 23), 55 at session C (SW = 31, SI = 24), and 47 at session D (SW = 23, SI = 24). For details on excluded subjects for each session, see Supplementary Table 1.

### 2.2 At home training

Figure 1A shows an overview of the study design and details on the training design can be found in Schwarze et al. (2025). Briefly, the two training groups differed only in the relative amount of task switching performed during each training session. More specifically, the SI group performed 83% single-tasking blocks and 17% task-switching blocks per training session and the SW group 17% single-tasking blocks and 83% task-switching blocks per training session. Children trained on a tablet at home for a total of 27 training sessions (30–40 min per session) with at least 24h between sessions and three training sessions per week. The rules and stimuli in each session were identical for both groups and all children performed the same number of trials on a given training session (i.e., differences in the duration of a training session occurred only due to different lengths of the self-paced breaks). Participants performed a task-switching paradigm in the MRI scanner (see details below) on four occasions throughout the training: before training (pre-test, session A), after ca. three weeks of training (session B), after ca. six weeks of training (session C), and once the training was completed after nine weeks (post-test, session D). The task-switching paradigm in the MRI scanner was identical to one of the trained paradigms.

**Figure 1:**
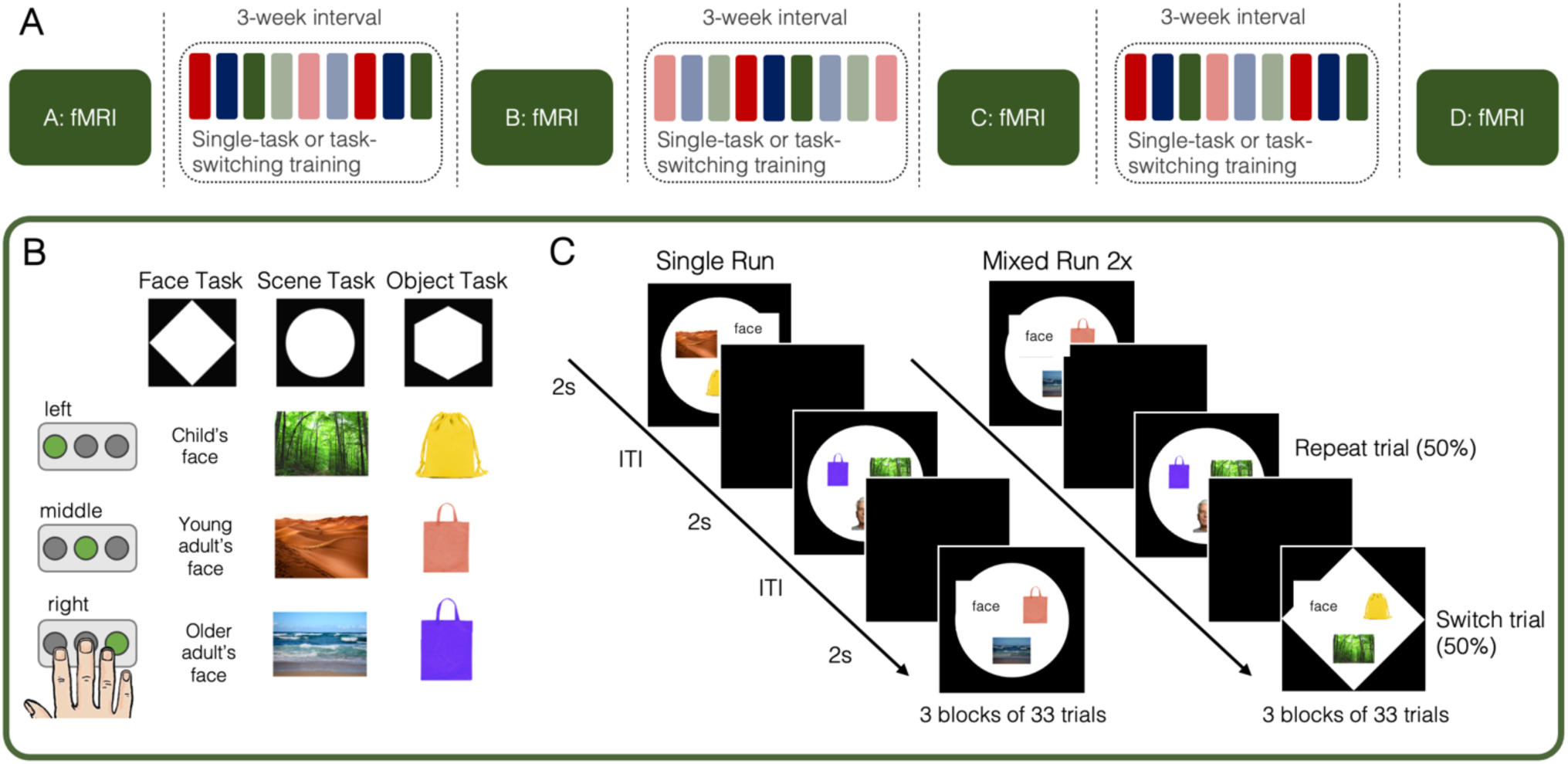
Study design and experimental task-switching paradigm. (A) Assessments and training across the nine weeks of the training. fMRI indicates that the main task-switching paradigm (see B and C) was performed in the MRI scanner. Colored bars indicate the training sessions. (B) The task-switching paradigm completed in the fMRI scanner. The shape cue indicated one of the three tasks. As indicated by three exemplar stimuli of each task, participants selected one of the three buttons based on the face’s age in the face task, the type of environment in the scene task, and the color of the object in the object task. (C) Three sequential trials of the single and mixed blocks; in the single block depicted here, participants performed the scene task on every trial. In the mixed task, the shape cues (and therefore tasks) repeated on some trials and switched on others. ITI: Inter-trial interval. Image credits: Young and old adult FACES were taken from the FACES collection (Ebner et al., 2010). Figure adapted from <u>Schwarze et al. (2025)</u>, under <u>CC.BY 4.0</u>.

### 2.3 Experimental task-switching paradigm

At each of the four MRI sessions, children completed a task-switching paradigm in the MRI scanner (see also Schwarze et al., 2023, for detailed paradigm description). They had familiarized themselves with the paradigm in an assessment session completed prior to the first MRI session, and during two practice blocks completed in an MRI simulator before performing the task in the MRI scanner. In the MRI scanner, participants watched a muted cartoon during the initial T1-weighted scan and subsequently performed three runs (99 trials each) of the experimental paradigm. The paradigm required participants to categorize stimuli (via button presses) based on three different rules that were indicated by the shape in the background (Figure 1B–C). Every trial lasted 2 s, followed by a fixation cross (1–6s, jittered) along with an extended fixation period (20 s) after every 33 trials. The three tasks were presented sequentially in the first run (i.e., single run), and intermixed (50% switch rate with unpredictable switches) in runs two and three (i.e., mixed runs). The first trial of each run was excluded from all behavioral analyses, as it could not be categorized as switch or repeat trial.

### 2.4 Behavioral analyses

In our previous work, we observed training-related improvements in accuracy in this sample (Schwarze et al., 2025). We therefore preregistered our hypotheses with accuracy as the outcome variable for predicting individuals’ training success from functional network modularity (see https://aspredicted.org/vh6x-q55f.pdf). Accuracy was defined as the proportion of correct responses relative to given responses and calculated separately for the three conditions (single trials, repeat trials, and switch trials).

In addition to the pre-registered analysis of accuracy, we sought to combine accuracy and response speed to capture performance in the task-switching paradigm more fully. To this end, we had preregistered analyses on performance measures derived from hierarchical drift diffusion modeling (Wiecki et al., 2013). However, due to methodological concerns regarding the appropriateness of drift diffusion models for task switching paradigms, especially paradigms with more than two tasks such as the present paradigm (Ratcliff and Frank, 2012; Ratcliff and McKoon, 2008), we ultimately did not adopt this approach. An alternative and more appropriate approach for this kind of data were linear integrated speed-accuracy scores (LISAS; Vandierendonck, 2018, 2017) which allowed us to account for both speed and accuracy. Since individuals differ in prioritizing accuracy or speed even if the same instructions are given (Heitz, 2014), LISAS might thus more sensitively capture individual differences in task-switching performance and change therein. See Supplementary Table 2 for an overview of the changes from the preregistration (based on Willroth and Atherton, 2024).

LISAS were calculated after trials with response times below 200 ms and above 3000 ms (i.e., responses given after stimulus presentation and the first second of the subsequent fixation cross), and trials with no responses were removed. Analyses were conducted using linear mixed models within a Bayesian framework (brms; Bürkner, 2017). To examine performance changes with training, the model of LISAS as the dependent variable included a fixed linear effect of session to capture linear change in performance and a fixed quadratic effect of session to capture a potential deceleration of the change in performance. Additional fixed effects included condition (single vs. repeat vs. switch) and group (SI vs. SW) and were modeled to interact with the linear and quadratic session effects. The model included a random intercept of the participant and random slopes of the session. Reported effects are based on 95% credible intervals (CI), meaning that we can make a statement with 95% probability (cf. Bürkner, 2017; see also Morey et al., 2016). For the corresponding model of accuracy see Schwarze et al. (2025) and summary of results below.

### 2.5 fMRI data acquisition and preprocessing

MR images (functional and structural) were collected on a 3-Tesla Siemens Tim Trio MRI scanner. Structural images (220 slices; 1 mm isotropic voxels; TR = 4500 ms; TE = 2.35 ms; FoV = 160 x 198 x 220) and three functional runs, each consisting of 230 whole-brain echo-planar images of 36 interleaved slices (TR = 2000 ms; TE = 30 ms; 3 mm isotropic voxels), were acquired at each of the four timepoints. Preprocessing was performed using fMRIprep (Version 20.2.0; Esteban et al., 2019). For a detailed description see the fMRIprep documentation (https://fmriprep.org/en/stable/). Briefly, functional images were co-registered to individual anatomical templates using FreeSurfer (Greve and Fischl, 2009). The anatomical template was created from anatomical scans across all sessions, removing scans that were of poor quality based on the MRIQC classifier (Version 0.15.2; Esteban et al., 2017) and additional visual inspection. Images were slice-time corrected (using AFNI; Cox and Hyde, 1997), realigned (using FSL 5.0.9; Jenkinson et al., 2002), and resampled into MNI152NLin6Asym standard space with an isotropic voxel size of 2 mm. Finally, images were spatially smoothed with an 8 mm FWHM isotropic Gaussian kernel using SPM12 (Functional Imaging Laboratory, University College London, UK).

### 2.6 fMRI data analysis

#### 2.6.1 Finite impulse response general linear models (FIR GLM)

Based on Cole et al. (2019), we regressed out the main effects of the task on the BOLD time series using finite impulse response general linear models (FIR GLM) before constructing the functional connectomes. Data were high-pass filtered at 128 s and the first five volumes of each run were discarded to allow for stabilization of the magnetic field. FIR GLM analyses were performed using SPM12 software (Functional Imaging Laboratory, UCL, UK). For each participant, we estimated an event-related GLM that modeled a separate impulse regressor to each 2 s interval within a 20 s window following each trial onset. These ten FIR regressors per trial were modeled separately for correct single, correct repeat, correct switch trials, and any excluded trials (i.e., incorrect trials, trials with no responses, and the first trial of each run). Framewise displacement per volume in mm (Power et al., 2012), realignment parameters (three translation and three rotation parameters), and the first six anatomical CompCor components (as provided by fMRIprep; Behzadi et al., 2007) were included as regressors of no interest to reduce the effect of head motion artifacts. Temporal autocorrelations were estimated using first-order autoregression.

#### 2.6.2 Connectivity matrices and network assignment

For each participant, we created a connectivity matrix based on the pairwise correlations of the residual time series in the 400 parcels of the Schaefer parcellation (Schaefer et al., 2018). In a graph-theoretical framework, each parcel denotes a node in the ensuing graph. Connections between nodes represent the edges, with their weight being defined by the correlation strength between the time series of the two nodes that a given edge connects. Each of the 400 parcels was assigned to one of seven functional subnetworks (visual, somatomotor, dorsal attention, ventral attention, limbic, frontoparietal, default mode) based on the child-specific network configuration of the Schaefer parcels by Tooley et al. (2022a). Before calculating modularity, we thresholded the graph (i.e., the correlation matrix) at five different thresholds (in 2% steps between 2 and 10%; cf. Baniqued et al., 2019), keeping only the strongest connections (i.e., the edges with the highest weights), and only including positive connections. All subsequent analysis steps were performed for each of these thresholds separately.

#### 2.6.3 Network modularity index

Modularity was calculated on unweighted connections (i.e., binarized connectivity matrices) at each threshold. We calculated modularity using the igraph package in R (Csárdi et al., 2024), based on the modularity definition by Clauset et al. (2004). Using this approach, we derived one network modularity index for each individual at each session, with higher values, i.e., approaching 1, indicating a more modular organization. Thus, higher modularity values describe a network configuration that, at a certain threshold of connectivity strength, exhibits more connections between nodes that belong to the same functional subnetwork than between nodes that belong to different functional subnetworks. There were no outliers in modularity indices for each threshold at any of the sessions, defined as participants with modularity indices of > 3.5 standard deviations from the mean (p < 0.001, two-tailed).

### 2.7 Statistical analysis of network modularity

We first tested whether functional networks became more modular with training. To this end, we used linear-mixed models and tested for changes in the modularity index of each session (as the dependent variable) by the fixed effects of training group (SI vs. SW) and session, including participants as random intercepts and random slopes of session. Model comparisons using leave-one-out cross-validation (Vehtari et al., 2022) suggested that across thresholds, the model without any interactions of group either fit better or did not differ from the model including interactions of group (see Supplementary Table 3). We therefore report the results of the models without interactions involving training group.

Next, we tested whether functional network modularity before training or changes in functional network modularity with training were associated with performance changes. To this end, we predicted session-specific accuracy and LISAS (in separate models as the dependent variables indexing behavioral performance) by children’s network modularity at the pre-training session as well as session-specific change in network modularity relative to the pre-training session. Specifically, models included fixed effects of the modularity index at session A, session-specific change in the modularity index (i.e., difference in the modularity indices of each session [B, C, and D] relative to session A), condition (single vs. repeat vs. switch), training group (SI vs. SW), and linear and quadratic effects of session, with random effects for participants and random slopes for session. To assess whether the effects on change in performance of modularity at session A or change in modularity differed between the two training groups, we compared a model that allowed for interactions with the group effect to a model that did not allow for interactions with the group effect. Leave-one-out cross validation (Vehtari et al., 2022) did not favor the more complex model including the interactions of the group effect (see Supplementary Table 4). Thus, we report the results of the simpler model without any interactions of the group effect. Additionally, to save degrees of freedom and to keep models focused on our hypotheses, models did not allow for any interactions between pre-training modularity and change in modularity, nor between the linear and quadratic effect of session.

Before the main analyses testing the association between modularity and performance change, we sought to replicate previously reported age differences in network modularity. Specifically, we tested whether pre-training modularity was associated with age among children using Pearson’s correlation. Additionally, we tested whether children at the pre-test session showed the expected pattern of less modular networks compared to the adult sample, using a paired t-test at each of the five thresholds. P-values were corrected for multiple comparisons using false-discovery rate (FDR) correction (Benjamini and Hochberg, 1995). Moreover, we tested whether modularity at the pre-training session was associated with performance (i.e., LISAS) at the same session, again using Bayesian linear mixed models of LISAS as dependent variable, modularity and condition as fixed effects, and participant as a random effect.

Finally, we explored how the contribution of each subnetwork to individuals’ pre-training modularity scores and change therein impacted the effect on training outcomes. To this end, we re-calculated each individual’s modularity index (at the 10% threshold) iteratively excluding a single subnetwork and then tested whether we still observed any effect of pre-training modularity or change in modularity on training outcomes. Since modularity captures the organization of a network consisting of multiple subnetworks, it cannot be meaningfully calculated for single subnetworks. Thus, to measure the impact of a specific subnetwork, we removed all connections involving this subnetwork and assessed whether the modularity index became a less meaningful predictor of training outcomes as a result by estimating the model testing the association of change in LISAS with modularity at session A and change in modularity described above (see Supplementary Tables 8 and 9, and Supplementary Figure 1).

## 3. Results

### 3.1 Improved performance with training

We have previously reported that the children in both training groups showed reduced accuracy mixing costs after the first training sessions, with this improvement slowing down towards the end of training (Schwarze et al., 2025). In contrast, switch costs did not change with training. Overall, the changes in LISAS reported here closely match the previously reported effects of training on performance. As a basis for the interpretation of the analyses of individual differences below, we briefly report changes in LISAS in the two training groups which are of interest here.

Changes in LISAS over the course of training for each condition are shown in Figure 2. Across sessions, LISAS were lowest on single trials, followed by repeat trials (single vs. repeat: est. = –0.33; 95%-Credible Interval (CI) –0.38, –0.28), and switch trials (switch vs. repeat: est. = 0.31; 95%-CI 0.25, 0.36), demonstrating that the paradigm elicited the expected mixing and switch costs. With training, performance improved, as evident in LISAS decreasing across conditions (linear effect of session: est. = –0.13; 95%-CI –0.19, –0.06). Improvements slowed down towards the end of training (quadratic effect of session: est. = 0.03; 95%-CI 0.01, 0.06). There was no evidence for any differences between groups or conditions in this change (i.e., the 95%-intervals bordered or included zero). Thus, performance improved in children undergoing single-task and task-switching training. Of note, we also observed substantial individual differences in the magnitude of these changes (i.e., random slope of linear effect of session: est. = 0.05; 95%-CI 0.03, 0.06). This raises the question of potential antecedents of individual differences in training-related improvements. Here, we examined the role of functional network organization for variability in training benefits. To contextualize any such findings, we began by testing for differences in network organization between children and adults.

**Figure 2:**
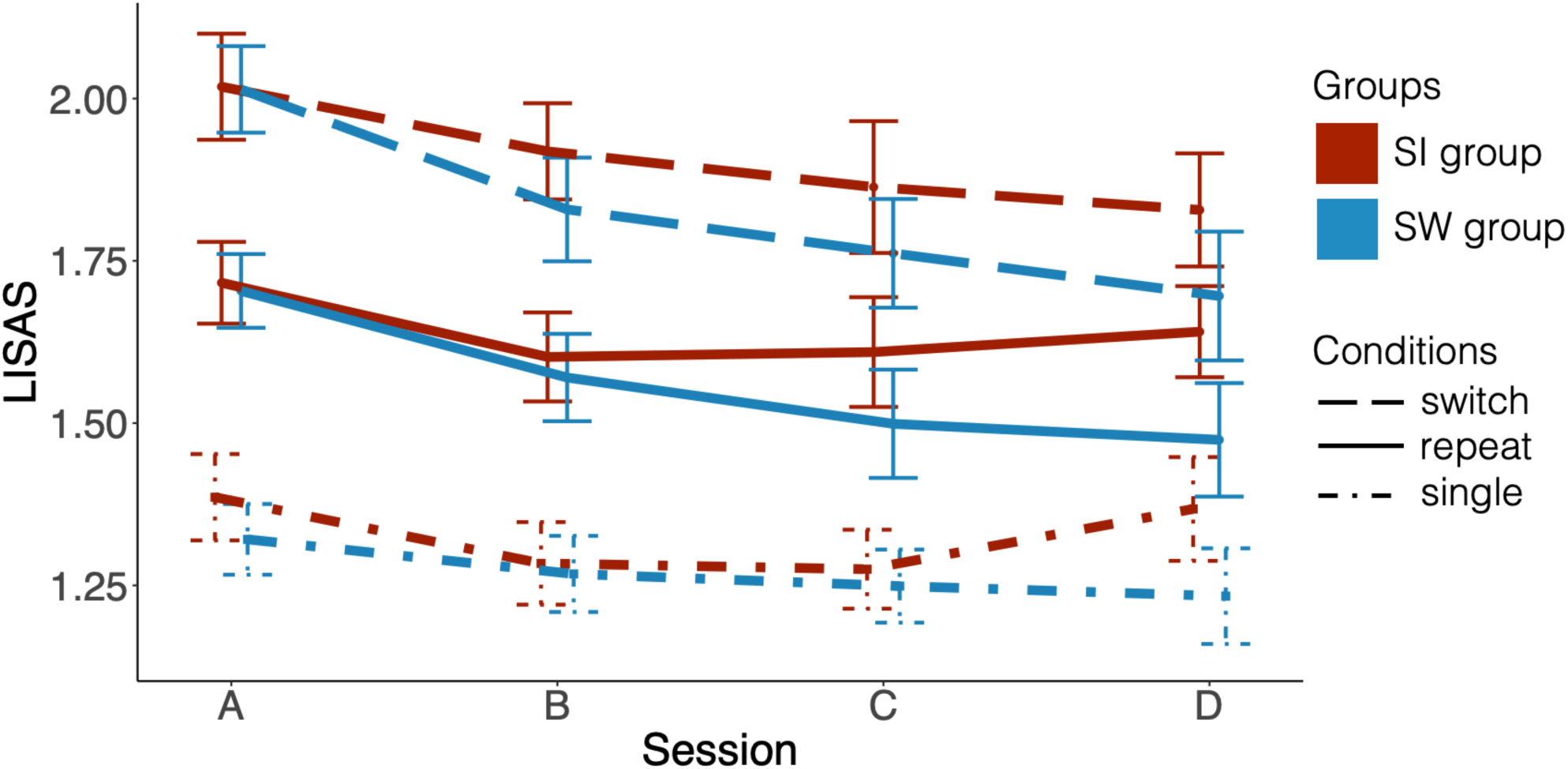
Changes in LISAS over the course of training. LISAS (lower scores indicate better performance) for each of the three conditions: switch trials (dashed line), repeat trials (solid line), and single trials (dot-dashed line). The SI group is shown in red, and the SW group in blue. Error bars denote 95%-confidence intervals.

### 3.2 Children show less modular network organization than adults

To test for the predicted pattern of less modular network organization in children than adults, we compared children’s modularity scores before they started training (i.e., at Session A) to the comparison group of adults. As shown in Figure 3A, modularity scores in children (M [across thresholds] = 0.33, SD [across thresholds] = 0.094) were significantly lower (p_FDR_ < .001 for each threshold, FDR-corrected for multiple comparisons) than in adults (M [across thresholds] = 0.42, SD [across thresholds] = 0.071). Within the group of children, modularity was not associated with age (mean r [across thresholds] = .019; all p_FDR_ > .84; Figure 3B). Finally, we did not observe any evidence for associations between modularity at the pre-training session and LISAS at the pre-training sessions (all 95%-CI included zero).

**Figure 3:**
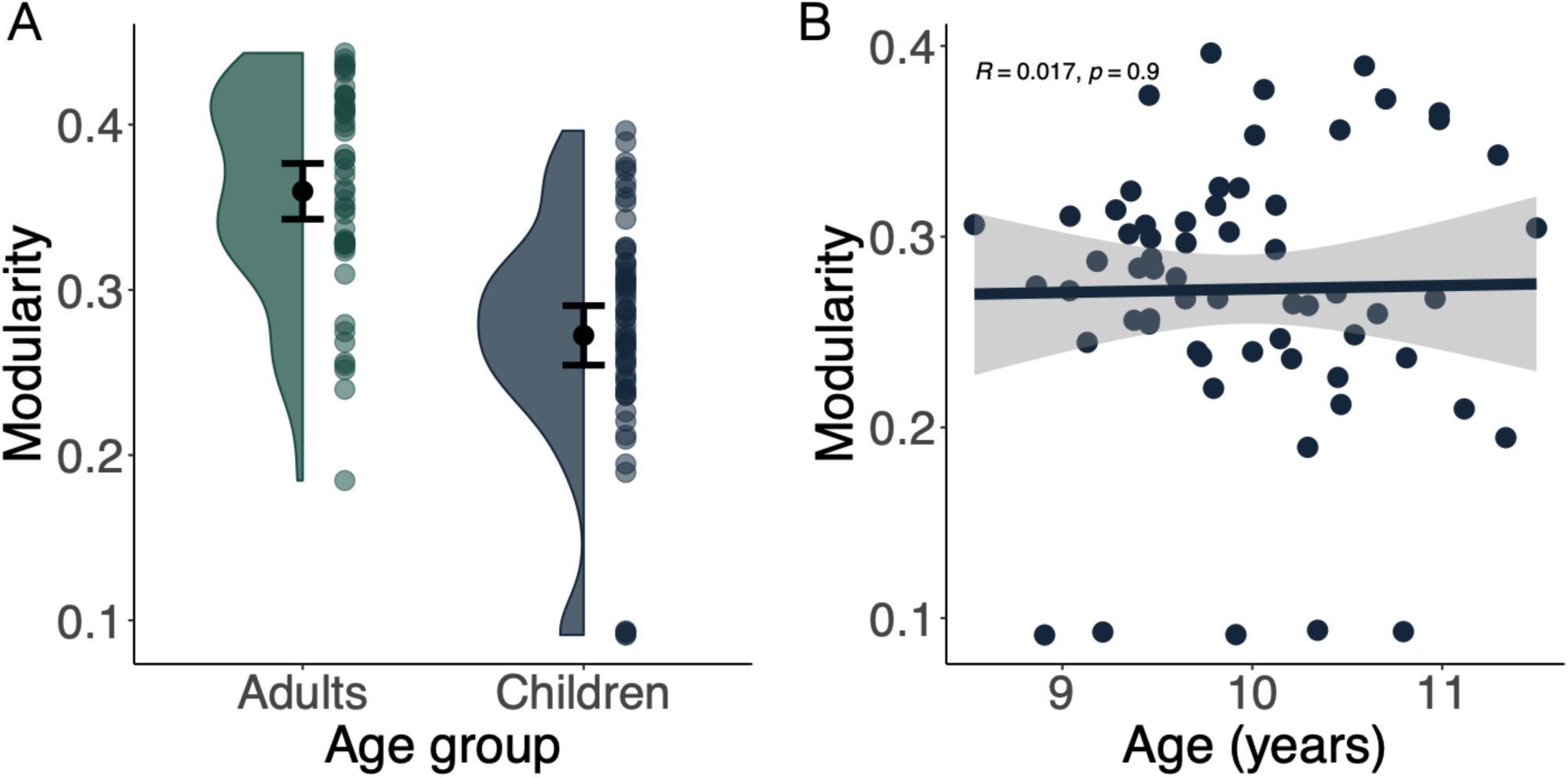
Age differences in modularity indices (at 10% threshold). (A) Modularity indices for adults (left) shown in green, and children (right) shown in blue. Error bars denote 95%-confidence intervals. (B) Correlation between modularity index and age in the group of children.

Taken together, this set of control analysis suggests that modularity captures a unique aspect of development independent of individuals’ age or pre-training performance.

### 3.3 Limited evidence for changes in network modularity over the course of training

Next, we tested whether network organization became more modular over the course of training. Again, we conducted these analyses at each of the five predefined thresholds. Evidence for changes in modularity was mixed. Specifically, we observed a linear increase of network modularity at the 2% (est. = 0.02; 95%-CI 0.01, 0.03) and 10% threshold (est. = 0.02; 95%-CI 0.01, 0.04; Figure 4A). This effect was not evident with 95% probability at the other thresholds, as the 95%-CI bordered zero (4%: est. = 0.02; 95%-CI: 0.00, 0.04; 6%: est. = 0.02; 95%-CI: 0.00, 0.04; 8%: est. = 0.02; 95%-CI 0.00, 0.04). A negative quadratic effect of session suggested that this linear increase slowed down towards the end of training, but again, the 95%-CI bordered (negative) zero at all thresholds (at each threshold: est. = –0.01; 95%-CI: –0.01, –0.00), cautioning the interpretation of this result. Complete model outputs for all thresholds are reported in Supplementary Table 5.

**Figure 4:**
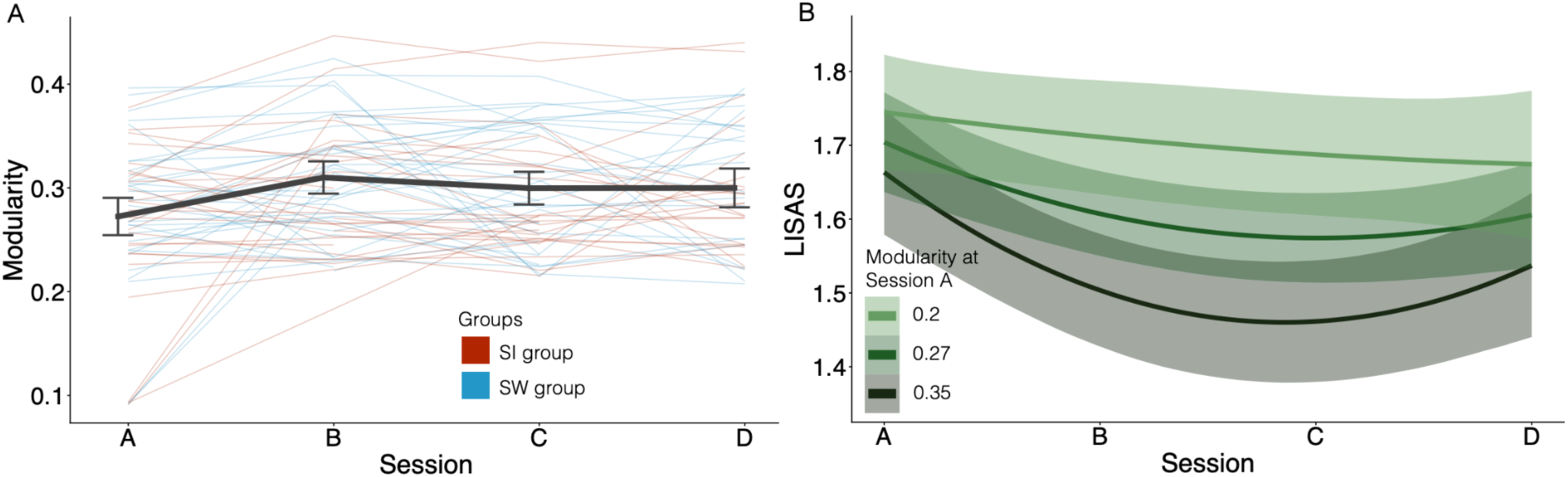
Changes in network modularity over the course of training and modularity at session A predicting changes in performance. (A) Changes in network modularity (10% threshold) across both groups. The thick line shows overall group mean; fine lines show individual changes in network modularity with individuals in the SI group shown in red, and individuals in the SW group shown in blue. Error bars denote 95%-confidence intervals. (B) Modularity (10% threshold) at session A predicting changes in LISAS with training. Estimated effects of the continuous modularity index are depicted at selected levels for visualization.

Taken together, we do not find convincing evidence that functional networks became more modular in organization over the course of training, or of any differences between the two training groups.

### 3.4 More modular network organization before training was associated with greater improvements in performance

Finally, we examined whether children who showed more modular networks before training, or greater changes in network modularity, showed greater improvements in task-switching performance with training. To this end, we tested for effects of session A modularity indices and for effects of the change in the modularity index on changes in the preregistered outcome variable of accuracy and in the exploratory outcome variable of LISAS (in separate models). Again, we conducted these analyses at all five thresholds. Since the results were consistent across thresholds, we only describe the effects for the model of the most lenient (i.e., 10%) threshold here; complete model outputs at all thresholds are reported in Supplementary Table 6 and 7.

For accuracy as the outcome measure, we observed a main effect of modularity at session A, indicating that children who showed higher modularity before training showed overall higher accuracy (est. = 0.55; 95%-CI: 0.09, 1.01). However, there was no evidence that pre-training network modularity or change in modularity over the course of training was associated with changes in accuracy over the course of training (all 95%-CI including zero).

When it comes to LISAS as the outcome measure, as shown in Figure 4B, we observed that higher modularity at session A was associated with greater decreases in LISAS across sessions (interaction of modularity at session A x linear effect of session: est. = –1.19; 95%-CI: –2.03, –0.36). Additionally, higher modularity at session A was associated with a more pronounced slowing down in performance improvements towards the end of training (interaction of modularity at session A x quadratic effect of session: est. = 0.36; 95%-CI: 0.07, 0.64). There were no effects of change in modularity on LISAS (all 95%-CI including zero), suggesting that changes in performance were not dependent on changes in network modularity with training. We did not observe any evidence for any condition-specific performance change dependent on modularity (all 95%-CI of interactions of conditions, modularity at session A and session included zero). Notably, with any of the subnetworks left out in control analyses, we continued to find evidence of pre-training modularity predicting performance changes with training, suggesting that the beneficial effect of modularity on training outcomes was driven by a whole-brain network consisting of all tested subnetworks (see Supplementary Table 8 for full statistics). Consistent with the computation of LISAS as an integrated measure of RT and accuracy, we observed a similar pattern of results with RT as dependent variable (see Supplementary Table 10). Finally, the association between modularity and change in LISAS remained when controlling for motion by including mean framewise displacement as a fixed effect (see Supplementary Table 11).

Taken together, the degree of modular organization of functional brain networks was positively associated with accuracy and with the rate of initial improvements with training in a combined measure of RTs and accuracy.

## 4. Discussion

The present study tested whether modularity as a marker of interindividual differences of functional brain network organization predicted improvements with task-switching training in a group of 8–11-year-old children who trained on task switching with varying intensity. Our previously published results of this sample (Schwarze et al., 2025) showed that, on average, accuracy improved in both training groups but that improvements were more pronounced in the task-switching training group relative to the passive control group. Building on these group comparisons in the present study, we asked whether individual differences in performance improvement with training were related to the modularity of functional brain networks. We found weak evidence overall for changes in network modularity with training. However, children who exhibited more modular organization of functional brain networks before the start of training showed overall higher accuracy and more pronounced, and faster improvements in a combined measure of RT and accuracy.

### 4.1 Network modularity predicting training outcomes

While children’s task-switching performance improved with training in both training groups, we observed substantial individual differences in training outcomes. Contrary to our hypotheses, network organization was only associated with overall levels of accuracy but not with change in accuracy. Further exploration of a performance score that combined children’s accuracy and speed of responding revealed that children with more modular networks prior to training showed more rapid improvements in task-switching performance with training. This pattern of results suggests that more modular functional networks allow for faster adaptation to task-switching demands and thus faster improvements with training, as operationalized by a score that more holistically captures performance across speed and accuracy. Strategy differences in prioritizing accuracy or speed (Heitz, 2014) might be further complicated in the context of training as individuals’ prioritization of accuracy or speed could change over the course of training (Heemskerk and Roebers, 2024). This could contribute to children with more modular network organization showing greater improvement in LISAS but not accuracy, as they might have learned more quickly how to balance speed and accuracy in a task-switching paradigm with training.

These results corroborate and extend previous findings with adults (Baniqued et al., 2019; Bassett et al., 2011; Ellefsen et al., 2015; Gallen et al., 2016) by demonstrating that individual differences in functional brain network organization were associated with training gains in children as well. There are multiple characteristics of modular networks that might put individuals with more modular organization in a position to benefit more from cognitive training. Biological networks are often large networks in which every connection is costly, such that modular organization may be particularly efficient (Tosh and McNally, 2015) and hence favored by evolution (Clune et al., 2013; Ellefsen et al., 2015). Increases in the efficiency of task processing, such as the speed in which single tasks or task components can be completed, are thought to at least partly contribute to training-related improvements in executive functions, including task switching (Dux et al., 2009; Kelly and Garavan, 2005; Poldrack, 2000; von Bastian et al., 2022). The speed in which single tasks or task components can be processed is especially relevant in the context of task switching, which requires the processing of multiple tasks in close temporal succession. Task-processing efficiency has been associated with better performance and greater training benefits in multitasking (Dux et al., 2009). Thus, individuals who show more modular network organization at the start of training might be particularly well prepared to meet the demands of the task-switching training which requires especially efficient task performance.

Additionally, modular networks show greater flexibility and adaptability to changing environments, potentially by allowing faster rearrangement of maladaptive or inefficient modules (Kashtan and Alon, 2005). Relatedly, modular networks support learning of different tasks or skills by preventing the forgetting of previously learned skills, presumably because different skills are processed in different modules (Ellefsen et al., 2015). Both these features, greater adaptability and reduced forgetting, could be particularly relevant in the present training regime, which included multiple task-switching paradigms differing in rule structure and stimuli (Schwarze et al., 2025). In the context of these multiple paradigms’ variations in rule structure, children in both training groups likely benefitted most from the training if they were able to keep the different rules and stimuli they encountered during training separate but could simultaneously generalize the overarching skills of maintaining, updating, and inhibiting rules as required during task switching. While research linking brain network modularity to learning multiple new tasks is scarce, artificial neural networks have been shown to generalize from different component tasks particularly well if they can be recombined flexibly, with generalization success supported by more modular network architecture (Lake and Baroni, 2023; see Cole, 2024, for a review).

Finally, even though we used a child-based network parcellation (Tooley et al., 2022a), we observed that adults showed greater network modularity than children, in line with previous observations that network organization of functional brain networks continues to develop over childhood (Baum et al., 2017; Fair et al., 2007; Keller et al., 2023; Luna et al., 2015; Sun et al., 2024; Wang et al., 2020; Yin et al., 2025). While network organization is still developing in children, more modular network organization of functional and structural brain networks has been associated with overall better cognitive performance (Baum et al., 2017; Wang et al., 2020), a pattern we also observed here with greater modularity associated with higher overall accuracy during task switching. Children with more modular networks may thus be already in a better initial position to learn the training paradigm and can more quickly draw benefits from the training compared to children with less modular networks, who showed slower training gain. One previous longitudinal study has explored the role of network modularity in the context of learning during child development: McCormick and colleagues (2021) showed a positive association between network modularity and feedback learning over the course of a single scan session of 20–30 minutes. However, this association only became apparent after adolescence and not in younger participants. By contrast, we found the association between modularity and training outcomes in a group of younger individuals between 8 and 11 years, perhaps because our participants completed an intensive task-switching or single-task training over nine weeks, and/or because of task differences. This contrast in results could suggest that the benefit of more modular networks requires greater training demands (as presented by the intensive task-switching and single-task training in the present study) in children, while it already becomes evident in short, less demanding learning tasks after adolescence (McCormick et al. (2021). To elucidate how the predictive nature of network modularity for training outcomes changes across development, it would therefore be important to combine longitudinal data of a wider age range with intensive cognitive training paradigms such as the one applied in the present study.

Taken together, our findings suggest that more modular functional brain networks position an individual in a more optimal state to meet increased task-switching demands of the intensive training regimen (Gallen and D’Esposito, 2019; Schultz and Cole, 2016) already in middle to late childhood, and thus earlier than previous studies exploring modularity during development (McCormick et al., 2021).

### 4.2 Lack of changes in network modularity over the course of training

Unlike previous studies (Bassett et al., 2015), we did not observe convincing evidence of changes in network modularity over the course of training in the present study. However, the task-switching training in the present study differed substantially from the study by Bassett and colleagues (2015), which examined motor skill training in relation to network modularity. Motor skill learning has been associated with increases in activation in the sensorimotor network and decreases in activation in control networks, along with decoupling of connections between sensorimotor and control regions (Dayan and Cohen, 2011; Floyer-Lea and Matthews, 2005; Grafton et al., 2002; Nguyen et al., 2025), suggesting an automatization of the motor skill with learning (Ackerman and Cianciolo, 2000; Fleishman, 1972). This decoupling or “autonomy” of the sensorimotor system also drove the increase in modularity observed in the previous study (Bassett et al., 2015).

While a modular network organization, specifically a segregated sensorimotor network, supports automatization of the motor skill by reducing interference from other subnetworks, training an executive functions task such as task switching requires networks to be flexibly reconfigured and integrated (Kupis et al., 2021; Nomi et al., 2017; Shine et al., 2016; Shine and Poldrack, 2018). Specifically, task switching relies on a widespread set of brain regions (Richter and Yeung, 2014; Schwarze et al., 2023; Worringer et al., 2019) among which connectivity has been shown to increase with training (Guerra-Carrillo et al., 2014; Jolles et al., 2013; Kundu et al., 2013; Mackey et al., 2013; Thompson et al., 2016). A more modular network configuration facilitates network reconfiguration (Rubinov et al., 2011; Sporns et al., 2000; Sporns and Betzel, 2016), which is crucial for task switching (Kupis et al., 2021), and thus might be beneficial to succeed in task-switching training. Thus, modular network organization might be beneficial for motor skill training because of a segregated sensorimotor network and for task-switching training because it facilitates flexible network reconfiguration. Note that these effects might differ between cognitive domains: Finc and colleagues (2020) showed increasing modularity with working memory training in adults (see also Finc et al., 2026 for a review). The different age groups across studies additionally raise the question of whether the extent of change in network modularity with training differs between age groups (cf. Iordan et al., 2021). Future studies contrasting motor skill learning and cognitive training across different domains and training intensities with regard to network changes are needed to further elucidate how these types of training differ in the changes they elicit in network organization across the lifespan.

### 4.3 Limitations and outlook

The present study is a first examination as to how individual differences in functional brain networks contribute to effects of cognitive training in childhood. While we demonstrated that network modularity already predicts training outcomes during the period in which functional networks are still developing, open questions remain that need to be investigated in future studies.

First, while more modular networks were beneficial for training gain especially in the early phase of training, this effect leveled off towards the end of training. This might be due to ceiling effects of performance. However, even at the end of training, children did not reach adult levels of performance in the present paradigm (cf. Schwarze et al., 2023, 2025), speaking against a ceiling effect of the paradigm itself. Nonetheless, this pattern suggests that we might have missed more fine-grained changes in performance towards the end of training. Particularly in children with more modular networks, who showed earlier training gains, the present training might not have been demanding enough to elicit further substantial improvements in performance (cf. Lövdén et al., 2010) towards the end of training, which we could have captured in the present paradigm. Future work should explore adaptive training paradigms to test whether children with more modular networks reach more demanding levels or whether the adaptive nature of such paradigms might reduce the effect of network modularity on training outcomes as it matches training demands to individuals’ capabilities and training needs.

Moreover, individual differences in modularity observed in children likely reflect a mixture between individual differences in the timing of modularity development and stable individual differences in modularity. These two sources of individual differences may contribute differentially to the observed association with training outcomes. Analyses of longitudinal data sets are needed to disentangle the two sources and examine their differential contributions to cognitive development.

Additionally, there are multiple ways to calculate modularity (Newman and Girvan, 2004; Sporns and Betzel, 2016). One main distinction is whether network affiliation of each node is predefined by a network parcellation or whether modularity is algorithmically maximized, resulting in a determination of network affiliation for different nodes for each individual. Both approaches have been used in developmental (Gu et al., 2020) and training contexts (Arnemann et al., 2015; Baniqued et al., 2019; Gallen et al., 2016). Here, we selected the former approach, as the latter approach of maximizing modularity within each participant would result in individual- and session-specific network parcellations, hindering the interpretation of training-related changes in modularity. Nonetheless, it is important to point out that individuals differ in the specific anatomical location of functional networks (e.g., Glasser et al., 2016; Gratton et al., 2018) with particularly pronounced individual differences in developmental populations (Cui et al., 2020; Keller et al., 2023, 2022). Future longitudinal studies should explore how the selection of modularity metrics and network parcellations influences the perceived developmental trajectory of network modularity.

Finally, we focused on network modularity due to its previous association with training outcomes. However, other metrics of network organization change over the course of childhood, e.g., network efficiency or hub organization (Demeter et al., 2023; Kolskår et al., 2018; Wu et al., 2013), and could impact training outcomes in addition to modularity. Future studies are needed to explore how different features of brain networks contribute to training outcomes across childhood and adolescence.

## Acknowledgments

The authors thank Julia Delius for editorial assistance and helpful comments.

## Funding

This work was supported by the Max Planck Institute for Human Development and by the DFG Priority Program SPP 1772 “Human performance under multiple cognitive task requirements: From basic mechanisms to optimized task scheduling” (Grant No. FA 1196/2-1 to Y.F.). Y.F.’s work was supported by a research fellowship from the Jacobs Foundation (2023-1510-00).

## Author contributions

Sina A. Schwarze (Data curation, Formal analysis, Investigation, Methodology, Visualization, Writing-original draft, Writing-review & editing), Ulman Lindenberger (Conceptualization, Methodology, Resources, Writing-review & editing), Silvia A. Bunge (Conceptualization, Methodology, Writing-review & editing) and Yana Fandakova (Conceptualization, Data curation, Funding acquisition, Investigation, Methodology, Project administration, Resources, Supervision, Writing-review & editing).

## Notes

### Competing Interest Statement

The authors have declared no competing interest.

### Summary of Updates

The introduction has been revised to improve the communication of the core research question. The methods section has been revised to clarify the training paradigm. The results section has been revised to clarify which analyses were preregistered. Supplementary files were updated.

