## Supplementary Materials for "Modular functional brain network organization contributes to training-related changes in task switching in children"

| **Content** | **Page** |
|---|---|
| *Supplementary Table 1:* Overview of excluded participants | 3 |
| *Supplementary Table 2:* Deviations from preregistration | 3 |
| *Supplementary Table 3:* Model comparisons for change in modularity | 3 |
| *Supplementary Table 4:* Model comparisons for association of LISAS and modularity across training | 4 |
| *Supplementary Table 5*: Complete model output for change in modularity across training | 4 |
| *Supplementary Table 6*: Complete model output for association between change in accuracy and modularity. | 4 |
| *Supplementary Table 7*: Complete model output for association between change in LISAS and modularity. | 5 |
| *Supplementary Table 8*: Complete model output for association between change in LISAS and modularity with one sub-network left out iteratively. | 5 |
| *Supplementary Table 9:* Complete model output for the differences in modularity index between sub-networks. | 6 |
| *Supplementary Figure 1:* Differences in modularity index at session A with different networks left out | 6 |
| *Supplementary Table 10*: Complete model output for association between change in response times (RT) and modularity. | 7 |
| *Supplementary Table 11*: Complete model output for association between change in LISAS and modularity controlling for head motion. | 7 |

*Supplementary Table 1:* Overview of excluded participants from each group for each session.

| | Session A | | Session B | | Session C | | Session D | |
|---|---|---|---|---|---|---|---|---|
| Group | SI | SW | SI | SW | SI | SW | SI | SW |
| Total | 44 | 43 | 44 | 43 | 44 | 43 | 44 | 43 |
| Too few training sessions | 1 | 1 | 1 | 1 | 1 | 1 | 1 | 1 |
| Dropout | 0 | 0 | 2 | 3 | 4 | 3 | 5 | 4 |
| Missing data due to scheduling issue | 0 | 0 | 1 | 0 | 0 | 0 | 0 | 0 |
| Missing data due to technical issues | 1 | 2 | 2 | 1 | 2 | 3 | 3 | 3 |
| Excluded due to poor performance or motion | 12 | 7 | 15 | 5 | 13 | 5 | 11 | 12 |
| Final included | 30 | 33 | 23 | 33 | 24 | 31 | 24 | 23 |

*Supplementary Table 2*: Deviations from preregistration (based on Willroth and Atherton, 2024). Overview of deviations from preregistration describing the type of deviation, the reason for deviation and the timing of the deviation. The "Reader Impact" column aims to position how the deviation impacts the interpretation of study results and their diagnosticity.

Deviations

| No. | Details | | Original wording | Deviation Description | Reader Impact |
|---|---|---|---|---|---|
| 1. | Type<br>Reasoning<br>Timing | Dependent variable<br>Researchers gained new knowledge<br>Before results known | The dependent variables of training-related changes in performance are average (i.e., averaged across single, repeat, and switch trials) accuracy (proportion correct of all given responses, see Schwarze et al., 2023) (...). Additionally, we will investigate the association of brain network modularity on training-related changes in mixing and switch costs (in accuracy, drift rate parameters, and boundary separation parameters) calculated as the difference between repeat and single-task trials, and switch and repeat trials, respectively. | Instead of running separate models on overall accuracy and accuracy costs, we ran models of condition-specific accuracy. In such a model, the main effect of modularity on accuracy would capture the effects on overall accuracy, while the interaction of modularity and condition would capture any effects on mixing or switch costs. We changed this after finding training-related improvements in accuracy mixing costs (Schwarze et al., 2025) that we wanted to accound for but before results of the present hypotheses were performed. | This deviation has a small impact on the reader, as all prergistered hypotheses are still testable and evident from this model. |
| 2. | Type<br>Reasoning<br>Timing | Dependent variable<br>Researchers gained new knowledge<br>Before results known | The dependent variables of training-related changes in performance are (...) and drift rate and boundary separation parameters derived from drift diffusion models (see Schwarze, Laube et al., 2023, bioRxiv). | Reviewers' on our previously published paper (Schwarze et al., 2025, the preprint cited in the preregistration), cautioned us agaist using drift diffusion models in the present paradigm. We therefore refrained from further analyzing them also in the present project. Searching for an alternative score that captured performance across both accuracy and speed, we came across linear integrated speed accuracy scores (Vandierendonck, 2017), which do not have the issues raised by the reviewers. We therefore applied these here as dependent variable in addition to the accuracy outcomes. | This deviation has medium impact on the reader. LISAS capture individuals performance on the task-switching paradigm, and the change therein, across speed and accuracy, thus allowing us to track changes in performance independent of potential strategy shifts in an individuals focus on accurate or fast responding. However, it does not hold information of the undelying cognitive processes and as such does cannot provide the same interpretation level as drift diffusion modeling. |

*Supplementary Table 3*: Model comparisons of the model testing for change in modularity with training showing that the model not including any interactions with group fit best or did not differ from the simpler model. Models were compared using the LOO function within the loo package in R (Vehtari et al. 2022). ELPD difference indicates the difference between each model's Bayesian leave-one-out estimate of the expected log pointwise predictive density (ELPD) and the best fitting model, whose ELPD difference is zero. SE indicates the standard error of this difference. Note that there is not a fixed ratio of SE to ELPD difference to indicate significant differences between models. Until these exist, developers' suggestions are that an ELPD difference around five times the SE can be interpreted as indicating differences between models. Bold values indicate the best fitting model.

| | Threshold | | | | | | | | | |
|---|---|---|---|---|---|---|---|---|---|---|
| | 10% | | 8% | | 6% | | 4% | | 2% | |
| Model | ELPD dif. | SE dif. | ELPD dif. | SE dif. | ELPD dif. | SE dif. | ELPD dif. | SE dif. | ELPD dif. | SE dif. |
| session + session(^2) + group | **0** | 0 | **0** | 0 | **0** | 0 | **0** | 0 | **0** | 0 |
| session * group + session(^2) * group | -2.0 | 1.0 | -1.6 | 1.3 | -1.7 | 1.2 | -1.6 | 1.3 | -2.3 | 1.4 |

*Supplementary Table 4*: Model comparisons of the models testing the association between change in LISAS and modularity (at session A and change in modularity) showing that the model not including any interactions with group fit best or did not differ from the simpler model.

| | Threshold | | | | | | | | | |
|---|---|---|---|---|---|---|---|---|---|---|
| | 10% | | 8% | | 6% | | 4% | | 2% | |
| Model | ELPD dif. | SE dif. | ELPD dif. | SE dif. | ELPD dif. | SE dif. | ELPD dif. | SE dif. | ELPD dif. | SE dif. |
| Model 1: no interaction of group factor | **0** | 0 | **0** | 0 | **0** | 0 | **0** | 0 | **0** | 0 |
| Model 2: interactions of group factor | -10.4 | 6.2 | -10.7 | 6.3 | -11.8 | 6.4 | -11.3 | 6.5 | -11.4 | 6.8 |

***Note***: Model 1 = mod_A * session * condition + mod_ch * session * condition + mod_A * session(^2) * condition + mod_ch * session(^2) * condition **+ group;** Model 2 = mod_A * session * condition *** group** + mod_ch * session * condition *** group** + mod_A * session(^2) * condition *** group** + mod_ch * session(^2) * condition * **group**

*Supplementary Table 5*: Complete model output for change in modularity across training in the two groups (SI vs. SW). CI indicates 95% credible intervals; bold values indicate estimates whose 95%-CI did not include zero.

| | Threshold | | | | | | | | | |
|---|---|---|---|---|---|---|---|---|---|---|
| Effect | 10% | | 8% | | 6% | | 4% | | 2% | |
| | Estimate | CI | Estimate | CI | Estimate | CI | Estimate | CI | Estimate | CI |
| Intercept | 0.28 | 0.26; 0.30 | 0.31 | 0.29; 0.33 | 0.34 | 0.32; 0.36 | 0.38 | 0.36; 0.40 | 0.42 | 0.41; 0.44 |
| session | **0.02** | 0.01; 0.04 | 0.02 | 0.00; 0.04 | 0.02 | 0.00; 0.04 | 0.02 | 0.00; 0.04 | **0.02** | 0.01; 0.03 |
| session(^2) | -0.01 | -0.01; -0.00 | -0.01 | -0.01; -0.00 | -0.01 | -0.01; -0.00 | -0.01 | -0.01; -0.00 | -0.01 | -0.01; -0.00 |
| group SW | 0.01 | -0.02; 0.03 | 0.01 | -0.02; 0.03 | 0.01 | -0.02; 0.03 | 0.01 | -0.02; 0.03 | 0.01 | -0.01; 0.03 |

*Supplementary Table 6*: Complete model output for association between change in accuracy and modularity. CI indicates 95% credible intervals; bold values indicate estimates whose 95%-CI did not include zero.

| | Threshold | | | | | | | | | | | | | | |
|---|---|---|---|---|---|---|---|---|---|---|---|---|---|---|---|
| Effect | 10% | | | 8% | | | 6% | | | 4% | | | 2% | | |
| | Estimate | CI | | Estimate | CI | | Estimate | CI | | Estimate | CI | | Estimate | CI | |
| Intercept | 0.62 | 0.49 | 0.75 | 0.62 | 0.49 | 0.75 | 0.61 | 0.47 | 0.75 | 0.60 | 0.46 | 0.75 | 0.62 | 0.46 | 0.78 |
| modularity at session A (mod_A) | **0.55** | 0.09 | 1.01 | **0.50** | 0.07 | 0.93 | **0.49** | 0.09 | 0.89 | **0.46** | 0.07 | 0.85 | 0.37 | -0.01 | 0.75 |
| session | 0.15 | -0.00 | 0.30 | 0.14 | -0.02 | 0.30 | 0.14 | -0.03 | 0.31 | 0.15 | -0.04 | 0.34 | 0.14 | -0.09 | 0.38 |
| condition (single vs. repeat) | **0.18** | 0.08 | 0.28 | **0.18** | 0.08 | 0.29 | **0.19** | 0.09 | 0.30 | **0.20** | 0.08 | 0.32 | **0.19** | 0.07 | 0.32 |
| condition (switch vs. repeat) | **-0.14** | -0.24 | -0.03 | **-0.14** | -0.24 | -0.03 | **-0.14** | -0.25 | -0.03 | **-0.15** | -0.26 | -0.03 | **-0.14** | -0.27 | -0.01 |
| change in modularity (mod_ch) | 0.69 | -0.19 | 1.60 | 0.65 | -0.16 | 1.50 | 0.67 | -0.10 | 1.48 | 0.64 | -0.14 | 1.44 | 0.60 | -0.19 | 1.41 |
| session(^2) | -0.05 | -0.10 | 0.01 | -0.04 | -0.10 | 0.01 | -0.04 | -0.10 | 0.02 | -0.04 | -0.11 | 0.02 | -0.04 | -0.13 | 0.04 |
| group (SW vs. SI) | **0.07** | 0.03 | 0.12 | **0.07** | 0.03 | 0.12 | **0.07** | 0.03 | 0.12 | **0.07** | 0.03 | 0.12 | **0.07** | 0.03 | 0.12 |
| mod_A * session | -0.21 | -0.71 | 0.30 | -0.17 | -0.65 | 0.31 | -0.16 | -0.63 | 0.31 | -0.16 | -0.64 | 0.32 | -0.14 | -0.67 | 0.40 |
| mod_A * condition single | -0.32 | -0.67 | 0.03 | -0.30 | -0.63 | 0.02 | **-0.31** | -0.63 | -0.01 | -0.30 | -0.62 | -0.00 | -0.26 | -0.56 | 0.03 |
| mod_A * conditionswitch | 0.21 | -0.16 | 0.57 | 0.21 | -0.13 | 0.54 | 0.20 | -0.13 | 0.51 | 0.19 | -0.12 | 0.50 | 0.15 | -0.16 | 0.45 |
| session* condition single | -0.02 | -0.22 | 0.17 | -0.02 | -0.22 | 0.19 | -0.02 | -0.24 | 0.21 | -0.02 | -0.27 | 0.23 | -0.00 | -0.30 | 0.30 |
| session * condition switch | 0.09 | -0.11 | 0.28 | 0.09 | -0.12 | 0.30 | 0.10 | -0.12 | 0.32 | 0.09 | -0.15 | 0.34 | 0.08 | -0.22 | 0.38 |
| mod_ch * session | -0.70 | -1.76 | 0.35 | -0.63 | -1.62 | 0.34 | -0.63 | -1.59 | 0.30 | -0.60 | -1.57 | 0.35 | -0.56 | -1.57 | 0.42 |
| mod_ch * condition single | -0.60 | -1.85 | 0.66 | -0.56 | -1.71 | 0.60 | -0.55 | -1.66 | 0.57 | -0.52 | -1.64 | 0.60 | -0.50 | -1.58 | 0.60 |
| mod_ch * condition switch | -0.05 | -1.32 | 1.21 | -0.05 | -1.23 | 1.12 | -0.09 | -1.22 | 1.02 | -0.10 | -1.22 | 1.00 | -0.13 | -1.25 | 0.97 |
| mod_A * session(^2) | 0.06 | -0.11 | 0.23 | 0.05 | -0.11 | 0.21 | 0.04 | -0.12 | 0.20 | 0.05 | -0.12 | 0.21 | 0.04 | -0.15 | 0.23 |
| condition single * session(^2) | 0.01 | -0.06 | 0.08 | 0.01 | -0.06 | 0.08 | 0.01 | -0.07 | 0.09 | 0.01 | -0.08 | 0.10 | 0.01 | -0.10 | 0.12 |
| condition switch * session(^2) | -0.03 | -0.09 | 0.04 | -0.03 | -0.10 | 0.04 | -0.03 | -0.11 | 0.04 | -0.03 | -0.12 | 0.05 | -0.03 | -0.14 | 0.08 |
| mod_ch * session(^2) | 0.22 | -0.06 | 0.51 | 0.20 | -0.07 | 0.47 | 0.20 | -0.06 | 0.46 | 0.19 | -0.08 | 0.46 | 0.17 | -0.11 | 0.46 |
| mod_A * session * condition single | -0.03 | -0.68 | 0.62 | -0.04 | -0.66 | 0.57 | -0.03 | -0.66 | 0.58 | -0.03 | -0.66 | 0.60 | -0.07 | -0.75 | 0.61 |
| mod_A * session * condition switch | -0.22 | -0.86 | 0.44 | -0.20 | -0.84 | 0.42 | -0.20 | -0.81 | 0.41 | -0.18 | -0.80 | 0.44 | -0.14 | -0.82 | 0.56 |
| mod_ch * session * condition single | 0.46 | -1.00 | 1.92 | 0.40 | -0.96 | 1.77 | 0.39 | -0.94 | 1.70 | 0.36 | -0.98 | 1.70 | 0.33 | -1.04 | 1.67 |
| mod_ch * session * condition switch | 0.15 | -1.31 | 1.60 | 0.14 | -1.23 | 1.51 | 0.17 | -1.13 | 1.48 | 0.18 | -1.13 | 1.52 | 0.24 | -1.13 | 1.61 |
| mod_A * session(^2) * condition single | -0.00 | -0.22 | 0.22 | 0.00 | -0.21 | 0.22 | 0.00 | -0.21 | 0.22 | -0.00 | -0.22 | 0.22 | 0.00 | -0.24 | 0.25 |
| mod_A * session(^2) * condition switch | 0.08 | -0.14 | 0.30 | 0.07 | -0.14 | 0.29 | 0.08 | -0.13 | 0.29 | 0.07 | -0.15 | 0.29 | 0.07 | -0.18 | 0.31 |
| mod_ch * session(^2) * condition single | -0.14 | -0.53 | 0.25 | -0.12 | -0.49 | 0.25 | -0.12 | -0.48 | 0.24 | -0.12 | -0.48 | 0.25 | -0.11 | -0.49 | 0.27 |
| mod_ch * session(^2) * condition switch | -0.03 | -0.41 | 0.36 | -0.02 | -0.39 | 0.34 | -0.02 | -0.37 | 0.33 | -0.03 | -0.39 | 0.34 | -0.04 | -0.42 | 0.34 |

*Supplementary Table 7*: Complete model output for association between change in LISAS and modularity. CI indicates 95% credible intervals; bold values indicate estimates whose 95%-CI did not include zero.

| Effect | 10% | | | 8% | | | 6% | | | 4% | | | 2% | | |
|---|---|---|---|---|---|---|---|---|---|---|---|---|---|---|---|
| | Estimate | CI | | Estimate | CI | | Estimate | CI | | Estimate | CI | | Estimate | CI | |
| Intercept | 1.88 | 1.70 | 2.05 | 1.87 | 1.69 | 2.04 | 1.87 | 1.69 | 2.05 | 1.87 | 1.67 | 2.06 | 1.86 | 1.65 | 2.06 |
| modularity at session A (mod_A) | -0.54 | -1.14 | 0.08 | -0.46 | -1.02 | 0.12 | -0.43 | -0.95 | 0.10 | -0.38 | -0.90 | 0.14 | -0.31 | -0.80 | 0.19 |
| session | 0.19 | -0.06 | 0.44 | 0.22 | -0.04 | 0.49 | 0.27 | -0.00 | 0.54 | **0.33** | 0.01 | 0.65 | **0.44** | 0.05 | 0.84 |
| condition (single vs. repeat) | **-0.38** | -0.52 | -0.23 | **-0.38** | -0.52 | -0.23 | **-0.38** | -0.53 | -0.23 | **-0.38** | -0.54 | -0.23 | **-0.38** | -0.55 | -0.21 |
| condition (switch vs. repeat) | **0.27** | 0.12 | 0.41 | **0.28** | 0.13 | 0.42 | **0.28** | 0.13 | 0.43 | **0.29** | 0.13 | 0.45 | **0.29** | 0.12 | 0.47 |
| change in modularity (mod_ch) | -1.15 | -2.78 | 0.49 | -1.03 | -2.54 | 0.46 | -1.00 | -2.43 | 0.40 | -0.82 | -2.19 | 0.55 | -0.88 | -2.24 | 0.46 |
| session(^2) | -0.07 | -0.15 | 0.02 | -0.08 | -0.17 | 0.02 | -0.09 | -0.19 | 0.00 | -0.11 | -0.22 | -0.00 | **-0.15** | -0.30 | -0.01 |
| group (SW vs. SI) | -0.04 | -0.11 | 0.03 | -0.04 | -0.11 | 0.03 | -0.04 | -0.11 | 0.03 | -0.04 | -0.11 | 0.03 | -0.04 | -0.11 | 0.03 |
| mod_A * session | **-1.19** | -2.03 | -0.36 | **-1.20** | -2.02 | -0.40 | **-1.23** | -1.99 | -0.46 | **-1.28** | -2.07 | -0.47 | **-1.39** | -2.30 | -0.50 |
| mod_A * condition single | 0.07 | -0.45 | 0.58 | 0.07 | -0.41 | 0.54 | 0.07 | -0.38 | 0.52 | 0.06 | -0.36 | 0.50 | 0.06 | -0.35 | 0.47 |
| mod_A * conditionswitch | 0.15 | -0.37 | 0.68 | 0.10 | -0.38 | 0.58 | 0.08 | -0.37 | 0.54 | 0.04 | -0.40 | 0.49 | 0.04 | -0.38 | 0.45 |
| session* condition single | -0.20 | -0.53 | 0.12 | -0.22 | -0.57 | 0.12 | -0.25 | -0.60 | 0.11 | -0.28 | -0.70 | 0.14 | -0.32 | -0.84 | 0.19 |
| session * condition switch | -0.05 | -0.38 | 0.28 | -0.06 | -0.41 | 0.28 | -0.05 | -0.42 | 0.31 | -0.05 | -0.46 | 0.37 | -0.01 | -0.53 | 0.50 |
| session * mod_ch | 0.31 | -1.66 | 2.27 | 0.18 | -1.62 | 2.00 | 0.12 | -1.58 | 1.85 | -0.11 | -1.85 | 1.60 | -0.18 | -1.90 | 1.60 |
| condition single * mod_ch | 0.94 | -1.29 | 3.16 | 0.91 | -1.11 | 2.94 | 0.89 | -1.03 | 2.85 | 0.77 | -1.10 | 2.66 | 0.92 | -0.89 | 2.74 |
| condition switch * mod_ch | 0.42 | -1.89 | 2.70 | 0.37 | -1.73 | 2.48 | 0.44 | -1.53 | 2.44 | 0.41 | -1.55 | 2.34 | 0.45 | -1.42 | 2.33 |
| mod_A * session(^2) | **0.36** | 0.07 | 0.64 | **0.36** | 0.09 | 0.64 | **0.37** | 0.11 | 0.64 | **0.39** | 0.11 | 0.67 | **0.45** | 0.13 | 0.77 |
| condition single * session(^2) | 0.09 | -0.03 | 0.20 | 0.10 | -0.03 | 0.22 | 0.10 | -0.02 | 0.23 | 0.12 | -0.03 | 0.27 | 0.13 | -0.05 | 0.32 |
| condition switch * session(^2) | 0.02 | -0.10 | 0.13 | 0.02 | -0.10 | 0.14 | 0.02 | -0.11 | 0.15 | 0.02 | -0.13 | 0.17 | 0.02 | -0.17 | 0.20 |
| mod_ch * session(^2) | 0.00 | -0.52 | 0.53 | 0.04 | -0.46 | 0.53 | 0.06 | -0.41 | 0.52 | 0.11 | -0.36 | 0.59 | 0.16 | -0.33 | 0.64 |
| mod_A * session * condition single | 0.85 | -0.24 | 1.95 | 0.85 | -0.20 | 1.90 | 0.83 | -0.18 | 1.83 | 0.85 | -0.23 | 1.91 | 0.86 | -0.30 | 2.03 |
| mod_A * session * condition switch | 0.07 | -1.04 | 1.17 | 0.11 | -0.94 | 1.18 | 0.08 | -0.95 | 1.10 | 0.07 | -1.00 | 1.12 | -0.02 | -1.18 | 1.17 |
| session * condition single * mod_ch | -0.33 | -2.97 | 2.31 | -0.29 | -2.73 | 2.15 | -0.27 | -2.66 | 2.06 | -0.11 | -2.45 | 2.21 | -0.25 | -2.62 | 2.07 |
| session * condition switch * mod_ch | -0.43 | -3.13 | 2.28 | -0.36 | -2.88 | 2.14 | -0.46 | -2.86 | 1.90 | -0.48 | -2.85 | 1.92 | -0.64 | -3.06 | 1.73 |
| mod_A * condition single * session(^2) | -0.31 | -0.68 | 0.07 | -0.31 | -0.68 | 0.06 | -0.30 | -0.66 | 0.05 | -0.31 | -0.69 | 0.07 | -0.31 | -0.73 | 0.11 |
| mod_A * condition switch * session(^2) | -0.06 | -0.44 | 0.32 | -0.07 | -0.44 | 0.29 | -0.07 | -0.42 | 0.29 | -0.06 | -0.44 | 0.32 | -0.04 | -0.46 | 0.37 |
| condition single * mod_ch * session(^2) | -0.02 | -0.71 | 0.67 | -0.03 | -0.68 | 0.61 | -0.03 | -0.65 | 0.60 | -0.07 | -0.70 | 0.57 | -0.03 | -0.68 | 0.61 |
| condition switch * mod_ch * session(^2) | 0.09 | -0.62 | 0.79 | 0.06 | -0.60 | 0.73 | 0.08 | -0.55 | 0.72 | 0.09 | -0.56 | 0.73 | 0.13 | -0.52 | 0.80 |

*Supplementary Table 8*: Complete model outputs for association between change in LISAS and modularity calculated with one sub-network left out iteratively (10% threshold only). CI indicates 95% credible intervals; bold values indicate estimates whose 95%-CI did not include zero.

| Effect | Left-out sub-network | | | | | | | | | | | | | | | | | | | | |
|---|---|---|---|---|---|---|---|---|---|---|---|---|---|---|---|---|---|---|---|---|---|
| | FPN | | | DAT | | | DMN | | | SMM | | | VIS | | | VAT | | | LIM | | |
| | Estimate | CI | | Estimate | CI | | Estimate | CI | | Estimate | CI | | Estimate | CI | | Estimate | CI | | Estimate | CI | |
| Intercept | 1.87 | 1.69 | 2.04 | 1.91 | 1.71 | 2.12 | 1.85 | 1.68 | 2.03 | 1.86 | 1.65 | 2.07 | 1.89 | 1.73 | 2.05 | 1.87 | 1.69 | 2.04 | 1.86 | 1.69 | 2.04 |
| modularity at session A (mod_A) | -0.43 | -0.95 | 0.10 | -0.53 | -1.10 | 0.05 | -0.46 | -1.09 | 0.18 | -0.52 | -1.33 | 0.30 | **-0.72** | -1.42 | -0.04 | -0.48 | -1.07 | 0.11 | -0.45 | -1.03 | 0.13 |
| session | 0.18 | -0.07 | 0.42 | 0.27 | -0.02 | 0.55 | 0.25 | -0.04 | 0.55 | 0.26 | -0.00 | 0.53 | 0.10 | -0.12 | 0.32 | 0.19 | -0.05 | 0.44 | 0.18 | -0.06 | 0.43 |
| condition (single vs. repeat) | **-0.38** | -0.52 | -0.24 | **-0.37** | -0.54 | -0.20 | **-0.38** | -0.53 | -0.24 | **-0.35** | -0.52 | -0.18 | **-0.39** | -0.52 | -0.25 | **-0.38** | -0.52 | -0.23 | -0.38 | -0.52 | -0.23 |
| condition (switch vs. repeat) | **0.27** | 0.13 | 0.42 | **0.25** | 0.08 | 0.42 | **0.29** | 0.14 | 0.44 | **0.27** | 0.09 | 0.44 | 0.26 | 0.12 | 0.39 | **0.27** | 0.12 | 0.42 | **0.27** | 0.12 | 0.42 |
| change in modularity (mod_ch) | -1.20 | -2.59 | 0.21 | -0.84 | -2.32 | 0.64 | -1.66 | -3.41 | 0.11 | -0.31 | -2.43 | 1.77 | -1.64 | -3.39 | 0.15 | -1.07 | -2.61 | 0.47 | -1.20 | -2.68 | 0.28 |
| session(^2) | -0.06 | -0.15 | 0.02 | -0.09 | -0.19 | 0.01 | -0.09 | -0.19 | 0.02 | -0.08 | -0.17 | 0.01 | -0.04 | -0.12 | 0.04 | -0.06 | -0.15 | 0.02 | -0.06 | -0.15 | 0.02 |
| group (SW vs. SI) | -0.04 | -0.11 | 0.04 | -0.04 | -0.11 | 0.04 | -0.04 | -0.11 | 0.04 | -0.04 | -0.11 | 0.04 | -0.04 | -0.11 | 0.03 | -0.04 | -0.11 | 0.03 | -0.04 | -0.11 | 0.03 |
| mod_A * session | **-1.00** | -1.71 | -0.28 | **-1.17** | -1.94 | -0.39 | **-1.48** | -2.51 | -0.45 | **-1.63** | -2.63 | -0.64 | **-1.05** | -1.95 | -0.16 | **-1.14** | -1.93 | -0.36 | **-1.09** | -1.86 | -0.32 |
| mod_A * condition single | 0.07 | -0.37 | 0.51 | 0.03 | -0.44 | 0.51 | 0.09 | -0.45 | 0.64 | -0.04 | -0.70 | 0.62 | 0.12 | -0.47 | 0.71 | 0.06 | -0.43 | 0.55 | 0.06 | -0.42 | 0.53 |
| mod_A * conditionswitch | 0.11 | -0.34 | 0.56 | 0.16 | -0.33 | 0.66 | 0.07 | -0.48 | 0.62 | 0.17 | -0.52 | 0.86 | 0.23 | -0.37 | 0.82 | 0.13 | -0.37 | 0.62 | 0.12 | -0.37 | 0.61 |
| session* condition single | -0.20 | -0.52 | 0.12 | -0.28 | -0.65 | 0.10 | -0.26 | -0.65 | 0.13 | -0.26 | -0.60 | 0.08 | -0.12 | -0.40 | 0.17 | -0.19 | -0.52 | 0.12 | -0.19 | -0.52 | 0.13 |
| session * condition switch | -0.06 | -0.39 | 0.26 | -0.07 | -0.44 | 0.32 | -0.11 | -0.50 | 0.28 | -0.04 | -0.39 | 0.32 | -0.02 | -0.31 | 0.27 | -0.04 | -0.37 | 0.28 | -0.05 | -0.37 | 0.28 |
| session * mod_ch | 0.51 | -1.14 | 2.16 | -0.07 | -1.81 | 1.68 | 0.70 | -1.46 | 2.85 | -0.73 | -3.15 | 1.74 | 0.86 | -1.25 | 2.94 | 0.27 | -1.58 | 2.12 | 0.45 | -1.31 | 2.21 |
| condition single * mod_ch | 0.95 | -0.96 | 2.85 | 0.79 | -1.26 | 2.82 | 1.42 | -0.97 | 3.79 | 0.82 | -1.98 | 3.67 | 1.05 | -1.35 | 3.44 | 0.85 | -1.24 | 2.96 | 0.93 | -1.08 | 2.93 |
| condition switch * mod_ch | 0.58 | -1.39 | 2.53 | 0.38 | -1.69 | 2.48 | 0.37 | -2.07 | 2.78 | 0.02 | -2.86 | 2.95 | 0.57 | -1.88 | 3.03 | 0.37 | -1.75 | 2.56 | 0.37 | -1.72 | 2.44 |
| mod_A * session(^2) | **0.30** | 0.06 | 0.54 | **0.36** | 0.08 | 0.62 | **0.45** | 0.09 | 0.81 | **0.47** | 0.13 | 0.81 | **0.32** | 0.02 | 0.62 | **0.34** | 0.08 | 0.61 | **0.33** | 0.06 | 0.59 |
| condition single * session(^2) | 0.09 | -0.03 | 0.20 | 0.11 | -0.02 | 0.24 | 0.11 | -0.02 | 0.25 | 0.10 | -0.02 | 0.22 | 0.05 | -0.04 | 0.15 | 0.08 | -0.03 | 0.20 | 0.08 | -0.03 | 0.20 |
| condition switch * session(^2) | 0.02 | -0.09 | 0.13 | 0.02 | -0.12 | 0.15 | 0.04 | -0.10 | 0.18 | 0.01 | -0.11 | 0.13 | 0.01 | -0.09 | 0.11 | 0.01 | -0.10 | 0.13 | 0.02 | -0.10 | 0.13 |
| mod_ch * session(^2) | -0.06 | -0.51 | 0.39 | 0.10 | -0.37 | 0.56 | -0.08 | -0.69 | 0.53 | 0.26 | -0.39 | 0.89 | -0.13 | -0.68 | 0.43 | 0.01 | -0.49 | 0.50 | -0.04 | -0.52 | 0.43 |
| mod_A * session * condition single | 0.72 | -0.22 | 1.66 | 0.89 | -0.15 | 1.89 | 1.08 | -0.26 | 2.44 | 1.17 | -0.11 | 2.47 | 0.68 | -0.49 | 1.85 | 0.79 | -0.25 | 1.83 | 0.76 | -0.26 | 1.78 |
| mod_A * session * condition switch | 0.12 | -0.84 | 1.07 | 0.10 | -0.95 | 1.11 | 0.31 | -1.06 | 1.67 | 0.02 | -1.33 | 1.36 | -0.04 | -1.23 | 1.16 | 0.06 | -1.00 | 1.11 | 0.08 | -0.95 | 1.09 |
| session * condition single * mod_ch | -0.41 | -2.65 | 1.83 | -0.14 | -2.52 | 2.26 | -0.62 | -3.52 | 2.28 | -0.07 | -3.40 | 3.17 | -0.66 | -3.45 | 2.16 | -0.28 | -2.79 | 2.20 | -0.39 | -2.76 | 2.00 |
| session * condition switch * mod_ch | -0.64 | -2.94 | 1.66 | -0.30 | -2.74 | 2.13 | -0.35 | -3.28 | 2.57 | 0.12 | -3.27 | 3.43 | -0.70 | -3.58 | 2.21 | -0.41 | -3.01 | 2.16 | -0.43 | -2.84 | 2.01 |
| mod_A * condition single * session(^2) | -0.26 | -0.59 | 0.06 | -0.32 | -0.67 | 0.04 | -0.41 | -0.89 | 0.06 | -0.40 | -0.85 | 0.04 | -0.24 | -0.64 | 0.15 | -0.28 | -0.64 | 0.07 | -0.27 | -0.62 | 0.08 |
| mod_A * condition switch * session(^2) | -0.06 | -0.39 | 0.26 | -0.05 | -0.41 | 0.31 | -0.15 | -0.63 | 0.33 | -0.04 | -0.50 | 0.42 | -0.03 | -0.44 | 0.37 | -0.05 | -0.41 | 0.31 | -0.05 | -0.40 | 0.30 |
| condition single * mod_ch * session(^2 | 0.01 | -0.58 | 0.61 | -0.06 | -0.69 | 0.57 | -0.01 | -0.80 | 0.79 | -0.08 | -0.92 | 0.78 | 0.09 | -0.64 | 0.82 | -0.03 | -0.68 | 0.63 | 0.01 | -0.62 | 0.64 |
| condition switch * mod_ch * session(^2 | 0.15 | -0.46 | 0.76 | 0.06 | -0.57 | 0.70 | 0.05 | -0.75 | 0.86 | -0.04 | -0.90 | 0.82 | 0.16 | -0.60 | 0.90 | 0.08 | -0.59 | 0.76 | 0.10 | -0.54 | 0.74 |

*Supplementary Table 9:* Complete model output for the differences in modularity index between sub-networks. As noted in the main text, a sub-network's contribution to modularity is assessed by removing all connections to regions of this sub-network and then calculating the modularity index again. CI indicates 95% credible intervals; bold values indicate estimates whose 95%-CI did not include zero.

left-out network used as reference

| Effect | DAT | | | FPN | | | DMN | | | LIM | | |
|---|---|---|---|---|---|---|---|---|---|---|---|---|
| | Estimate | CI | | Estimate | CI | | Estimate | CI | | Estimate | CI | |
| Intercept | 0.34 | 0.31 | 0.37 | 0.31 | 0.28 | 0.34 | 0.26 | 0.24 | 0.29 | 0.29 | 0.26 | 0.31 |
| DAT | | | | **0.03** | 0.03 | 0.03 | **0.08** | 0.07 | 0.08 | **0.05** | 0.05 | 0.05 |
| FPN | **-0.03** | -0.03 | -0.03 | | | | **0.05** | 0.05 | 0.05 | **0.02** | 0.02 | 0.03 |
| DMN | **-0.08** | -0.08 | -0.07 | **-0.05** | -0.05 | -0.05 | | | | **-0.02** | -0.03 | -0.02 |
| LIM | **-0.05** | -0.05 | -0.05 | **-0.02** | -0.03 | -0.02 | **0.02** | 0.02 | 0.03 | | | |
| SMM | **-0.10** | -0.10 | -0.10 | **-0.07** | -0.07 | -0.07 | **-0.02** | -0.03 | -0.02 | **-0.05** | -0.05 | -0.04 |
| VAT | **-0.06** | -0.06 | -0.06 | **-0.03** | -0.03 | -0.03 | **0.02** | 0.01 | 0.02 | **-0.01** | -0.01 | -0.01 |
| VIS | **-0.13** | -0.13 | -0.13 | **-0.10** | -0.10 | -0.10 | **-0.05** | -0.06 | -0.05 | **-0.08** | -0.08 | -0.08 |
| group (SW vs. SI) | 0.01 | -0.03 | 0.05 | 0.01 | -0.03 | 0.05 | 0.01 | -0.03 | 0.05 | 0.01 | -0.03 | 0.05 |

| Effect | SMM | | | VAT | | | VIS | | |
|---|---|---|---|---|---|---|---|---|---|
| | Estimate | CI | | Estimate | CI | | Estimate | CI | |
| Intercept | 0.24 | 0.21 | 0.27 | **0.28** | 0.25 | 0.31 | **0.21** | 0.18 | 0.24 |
| DAT | **0.10** | 0.10 | 0.10 | **0.06** | 0.06 | 0.06 | **0.13** | 0.13 | 0.13 |
| FPN | **0.07** | 0.07 | 0.07 | **0.03** | 0.03 | 0.03 | **0.10** | 0.10 | 0.10 |
| DMN | **0.02** | 0.02 | 0.03 | **-0.02** | -0.02 | -0.01 | **0.05** | 0.05 | 0.06 |
| LIM | **0.05** | 0.04 | 0.05 | **0.01** | 0.01 | 0.01 | **0.08** | 0.08 | 0.08 |
| SMM | | | | **-0.04** | -0.04 | -0.04 | **0.03** | 0.03 | 0.03 |
| VAT | **0.04** | 0.04 | 0.04 | | | | **0.07** | 0.07 | 0.07 |
| VIS | **-0.03** | -0.03 | -0.03 | **-0.07** | -0.07 | -0.07 | | | |
| group (SW vs. SI) | 0.01 | -0.03 | 0.05 | 0.01 | -0.03 | 0.05 | 0.01 | -0.03 | 0.05 |

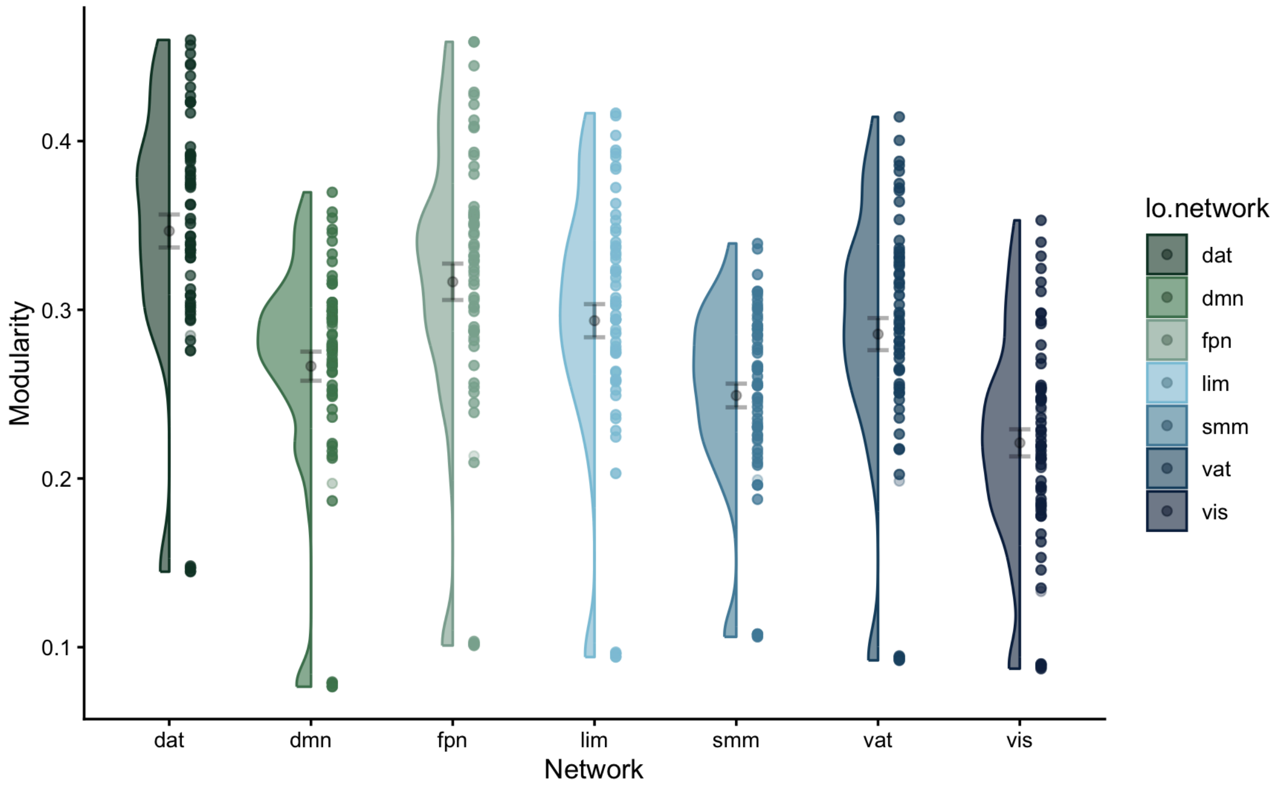


*Supplementary Figure 1: Differences in modularity index at session A with different networks left out (threshold = 10%; children only).* dat = Dorsal attention network, dmn = default mode network, fpn = frontoparietal network, lim = limbic network, smm = somatomotor network, vat = ventral attention network, vis = visual network

*Supplementary Table 10*: Complete model output for association between change in RT and modularity. CI indicates 95% credible intervals; bold values indicate estimates whose 95%-CI did not include zero.

| Effect | 10% | | | 8% | | | 6% | | | 4% | | | 2% | | |
|---|---|---|---|---|---|---|---|---|---|---|---|---|---|---|---|
| | Estimate | CI | | Estimate | CI | | Estimate | CI | | Estimate | CI | | Estimate | CI | |
| Intercept | 1.53 | 1.37 | 1.69 | 1.53 | 1.37 | 1.69 | 1.52 | 1.35 | 1.68 | 1.51 | 1.33 | 1.68 | 1.49 | 1.30 | 1.68 |
| modularity at session A (mod_A) | -0.23 | -0.79 | 0.33 | -0.19 | -0.70 | 0.33 | -0.14 | -0.63 | 0.34 | -0.10 | -0.58 | 0.37 | -0.05 | -0.50 | 0.39 |
| session | **0.23** | 0.02 | 0.45 | **0.26** | 0.04 | 0.48 | **0.30** | 0.06 | 0.54 | **0.36** | 0.09 | 0.63 | **0.47** | 0.14 | 0.79 |
| condition (single vs. repeat) | **-0.25** | -0.37 | -0.12 | **-0.25** | -0.37 | -0.12 | **-0.23** | -0.36 | -0.11 | **-0.23** | -0.37 | -0.09 | **-0.21** | -0.36 | -0.06 |
| condition (switch vs. repeat) | **0.20** | 0.07 | 0.33 | **0.21** | 0.08 | 0.34 | **0.22** | 0.09 | 0.36 | **0.23** | 0.08 | 0.37 | **0.23** | 0.08 | 0.39 |
| change in modularity (mod_ch) | -0.62 | -2.01 | 0.75 | -0.54 | -1.82 | 0.72 | -0.46 | -1.69 | 0.73 | -0.38 | -1.56 | 0.81 | -0.39 | -1.55 | 0.75 |
| session(^2) | **-0.08** | -0.15 | -0.01 | **-0.09** | -0.16 | -0.01 | **-0.10** | -0.18 | -0.02 | **-0.12** | -0.21 | -0.02 | **-0.16** | -0.27 | -0.04 |
| group (SW vs. SI) | -0.01 | -0.08 | 0.05 | -0.01 | -0.08 | 0.05 | -0.01 | -0.08 | 0.05 | -0.01 | -0.08 | 0.05 | -0.01 | -0.08 | 0.05 |
| mod_A * session | **-1.14** | -1.86 | -0.43 | **-1.14** | -1.83 | -0.46 | **-1.15** | -1.82 | -0.49 | **-1.19** | -1.88 | -0.52 | **-1.32** | -2.05 | -0.59 |
| mod_A * condition single | -0.10 | -0.55 | 0.34 | -0.09 | -0.50 | 0.32 | -0.12 | -0.51 | 0.26 | -0.12 | -0.49 | 0.24 | -0.15 | -0.51 | 0.20 |
| mod_A * conditionswitch | 0.17 | -0.30 | 0.63 | 0.13 | -0.30 | 0.55 | 0.09 | -0.32 | 0.49 | 0.05 | -0.34 | 0.45 | 0.04 | -0.34 | 0.41 |
| session* condition single | -0.21 | -0.48 | 0.05 | -0.23 | -0.51 | 0.04 | -0.26 | -0.56 | 0.03 | -0.29 | -0.63 | 0.04 | -0.36 | -0.77 | 0.04 |
| session * condition switch | -0.02 | -0.30 | 0.26 | -0.03 | -0.32 | 0.26 | -0.01 | -0.33 | 0.30 | 0.02 | -0.33 | 0.37 | 0.09 | -0.34 | 0.52 |
| session * mod_ch | -0.14 | -1.79 | 1.53 | -0.22 | -1.76 | 1.35 | -0.31 | -1.79 | 1.21 | -0.41 | -1.91 | 1.10 | -0.51 | -1.97 | 0.97 |
| condition single * mod_ch | 0.25 | -1.58 | 2.11 | 0.28 | -1.42 | 1.98 | 0.23 | -1.38 | 1.87 | 0.19 | -1.41 | 1.79 | 0.29 | -1.21 | 1.84 |
| condition switch * mod_ch | 0.19 | -1.66 | 2.08 | 0.14 | -1.60 | 1.89 | 0.15 | -1.49 | 1.83 | 0.25 | -1.38 | 1.90 | 0.38 | -1.17 | 1.97 |
| mod_A * session(^2) | **0.33** | 0.10 | 0.57 | **0.33** | 0.11 | 0.57 | **0.34** | 0.11 | 0.57 | **0.35** | 0.12 | 0.59 | **0.41** | 0.15 | 0.67 |
| condition single * session(^2) | 0.09 | 0.00 | 0.19 | 0.10 | 0.00 | 0.20 | 0.11 | 0.00 | 0.22 | 0.12 | 0.00 | 0.24 | 0.14 | -0.00 | 0.29 |
| condition switch * session(^2) | 0.01 | -0.09 | 0.10 | 0.01 | -0.09 | 0.11 | 0.00 | -0.11 | 0.12 | -0.01 | -0.13 | 0.12 | -0.03 | -0.18 | 0.13 |
| mod_ch * session(^2) | 0.16 | -0.28 | 0.61 | 0.18 | -0.24 | 0.59 | 0.20 | -0.22 | 0.60 | 0.22 | -0.19 | 0.63 | 0.27 | -0.14 | 0.68 |
| mod_A * session * condition single | 0.82 | -0.08 | 1.72 | 0.82 | -0.03 | 1.69 | 0.83 | -0.01 | 1.68 | 0.84 | -0.02 | 1.70 | 0.91 | -0.02 | 1.84 |
| mod_A * session * condition switch | -0.05 | -0.98 | 0.88 | -0.01 | -0.90 | 0.88 | -0.06 | -0.94 | 0.82 | -0.12 | -1.03 | 0.76 | -0.26 | -1.24 | 0.70 |
| session * condition single * mod_ch | 0.35 | -1.85 | 2.56 | 0.33 | -1.72 | 2.39 | 0.37 | -1.65 | 2.34 | 0.43 | -1.56 | 2.41 | 0.35 | -1.61 | 2.28 |
| session * condition switch * mod_ch | -0.20 | -2.47 | 2.01 | -0.13 | -2.25 | 1.97 | -0.22 | -2.28 | 1.79 | -0.42 | -2.48 | 1.62 | -0.72 | -2.74 | 1.28 |
| mod_A * condition single * session(^2) | -0.31 | -0.62 | 0.00 | -0.31 | -0.61 | -0.01 | **-0.30** | -0.60 | -0.01 | **-0.31** | -0.61 | -0.01 | -0.33 | -0.66 | -0.00 |
| mod_A * condition switch * session(^2) | -0.01 | -0.33 | 0.31 | -0.02 | -0.33 | 0.29 | 0.00 | -0.31 | 0.31 | 0.03 | -0.29 | 0.34 | 0.07 | -0.28 | 0.41 |
| condition single * mod_ch * session(^2) | -0.23 | -0.81 | 0.35 | -0.22 | -0.77 | 0.32 | -0.22 | -0.75 | 0.31 | -0.23 | -0.77 | 0.30 | -0.21 | -0.75 | 0.32 |
| condition switch * mod_ch * session(^2) | 0.03 | -0.56 | 0.62 | 0.01 | -0.55 | 0.57 | 0.04 | -0.50 | 0.59 | 0.10 | -0.45 | 0.65 | 0.18 | -0.37 | 0.73 |

*Supplementary Table 11*: Complete model output for association between change in LISAS and modularity controlling for head motion (i.e., mean framewise displacement). CI indicates 95% credible intervals; bold values indicate estimates whose 95%-CI did not include zero.

| Effect | 10% | | | 8% | | | 6% | | | 4% | | | 2% | | |
|---|---|---|---|---|---|---|---|---|---|---|---|---|---|---|---|
| | Estimate | CI | | Estimate | CI | | Estimate | CI | | Estimate | CI | | Estimate | CI | |
| Intercept | 1.84 | 1.65 | 2.02 | 1.83 | 1.65 | 2.02 | 1.83 | 1.65 | 2.01 | 1.84 | 1.64 | 2.04 | 1.82 | 1.60 | 2.02 |
| modularity at session A (mod_A) | -0.48 | -1.11 | 0.13 | -0.44 | -1.01 | 0.13 | -0.40 | -0.94 | 0.13 | -0.37 | -0.88 | 0.15 | -0.29 | -0.77 | 0.21 |
| session | 0.19 | -0.06 | 0.43 | 0.21 | -0.04 | 0.48 | 0.26 | -0.01 | 0.55 | 0.31 | 0.00 | 0.61 | **0.42** | 0.04 | 0.81 |
| condition (single vs. repeat) | **-0.38** | -0.52 | -0.23 | **-0.38** | -0.53 | -0.23 | **-0.38** | -0.53 | -0.23 | **-0.39** | -0.55 | -0.23 | **-0.39** | -0.55 | -0.22 |
| condition (switch vs. repeat) | **0.27** | 0.12 | 0.41 | **0.27** | 0.12 | 0.42 | **0.28** | 0.13 | 0.44 | **0.29** | 0.13 | 0.45 | **0.29** | 0.12 | 0.46 |
| change in modularity (mod_ch) | -1.05 | -2.67 | 0.59 | -0.95 | -2.47 | 0.59 | -0.91 | -2.32 | 0.53 | -0.78 | -2.15 | 0.58 | -0.82 | -2.13 | 0.46 |
| session(^2) | -0.07 | -0.15 | 0.02 | -0.08 | -0.17 | 0.01 | -0.09 | -0.19 | 0.00 | -0.11 | -0.22 | -0.00 | **-0.15** | -0.30 | -0.02 |
| group (SW vs. SI) | -0.04 | -0.12 | 0.03 | -0.04 | -0.11 | 0.03 | -0.04 | -0.11 | 0.03 | -0.04 | -0.12 | 0.03 | -0.04 | -0.11 | 0.03 |
| framewise displacement | 0.11 | -0.03 | 0.24 | 0.11 | -0.03 | 0.25 | 0.11 | -0.03 | 0.25 | 0.11 | -0.02 | 0.25 | 0.12 | -0.01 | 0.26 |
| mod_A * session | **-1.17** | -1.99 | -0.35 | **-1.16** | -1.96 | -0.38 | **-1.20** | -2.00 | -0.43 | **-1.21** | -1.97 | -0.42 | **-1.32** | -2.23 | -0.46 |
| mod_A * condition single | 0.06 | -0.45 | 0.58 | 0.07 | -0.41 | 0.55 | 0.06 | -0.40 | 0.50 | 0.07 | -0.35 | 0.51 | 0.07 | -0.34 | 0.47 |
| mod_A * conditionswitch | 0.15 | -0.37 | 0.68 | 0.11 | -0.37 | 0.60 | 0.08 | -0.38 | 0.53 | 0.05 | -0.37 | 0.49 | 0.04 | -0.36 | 0.46 |
| session * condition single | -0.21 | -0.54 | 0.11 | -0.23 | -0.57 | 0.11 | -0.25 | -0.62 | 0.11 | -0.28 | -0.66 | 0.12 | -0.33 | -0.82 | 0.15 |
| session * condition switch | -0.06 | -0.38 | 0.27 | -0.06 | -0.40 | 0.28 | -0.06 | -0.43 | 0.30 | -0.04 | -0.45 | 0.36 | -0.02 | -0.53 | 0.49 |
| session * mod_ch | 0.26 | -1.68 | 2.17 | 0.16 | -1.70 | 2.01 | 0.07 | -1.70 | 1.79 | -0.08 | -1.77 | 1.61 | -0.17 | -1.85 | 1.55 |
| condition single * mod_ch | 0.91 | -1.31 | 3.10 | 0.91 | -1.18 | 2.94 | 0.86 | -1.05 | 2.76 | 0.79 | -1.08 | 2.69 | 0.90 | -0.83 | 2.67 |
| condition switch * mod_ch | 0.41 | -1.84 | 2.68 | 0.40 | -1.72 | 2.52 | 0.43 | -1.58 | 2.43 | 0.43 | -1.52 | 2.38 | 0.45 | -1.38 | 2.31 |
| mod_A * session(^2) | **0.36** | 0.08 | 0.64 | **0.36** | 0.09 | 0.63 | **0.38** | 0.11 | 0.65 | **0.38** | 0.11 | 0.65 | **0.44** | 0.13 | 0.76 |
| condition single * session(^2) | 0.09 | -0.02 | 0.20 | 0.10 | -0.02 | 0.22 | 0.11 | -0.02 | 0.24 | 0.11 | -0.03 | 0.25 | 0.13 | -0.04 | 0.32 |
| condition switch * session(^2) | 0.02 | -0.09 | 0.13 | 0.02 | -0.10 | 0.14 | 0.03 | -0.10 | 0.16 | 0.02 | -0.12 | 0.16 | 0.02 | -0.17 | 0.20 |
| mod_ch * session(^2) | 0.02 | -0.50 | 0.54 | 0.05 | -0.45 | 0.55 | 0.07 | -0.39 | 0.55 | 0.11 | -0.34 | 0.57 | 0.17 | -0.31 | 0.64 |
| mod_A * session * condition single | 0.87 | -0.22 | 1.97 | 0.85 | -0.18 | 1.91 | 0.86 | -0.17 | 1.90 | 0.83 | -0.18 | 1.81 | 0.88 | -0.22 | 2.00 |
| mod_A * session * condition switch | 0.10 | -1.00 | 1.21 | 0.11 | -0.95 | 1.15 | 0.10 | -0.92 | 1.15 | 0.05 | -0.98 | 1.07 | -0.01 | -1.16 | 1.17 |
| session * condition single * mod_ch | -0.29 | -2.92 | 2.31 | -0.27 | -2.69 | 2.23 | -0.22 | -2.53 | 2.12 | -0.14 | -2.45 | 2.12 | -0.20 | -2.49 | 2.07 |
| session * condition switch * mod_ch | -0.41 | -3.10 | 2.26 | -0.39 | -2.91 | 2.12 | -0.44 | -2.82 | 2.01 | -0.52 | -2.87 | 1.79 | -0.63 | -3.01 | 1.81 |
| mod_A * condition single * session(^2) | -0.32 | -0.69 | 0.06 | -0.31 | -0.68 | 0.05 | -0.31 | -0.68 | 0.04 | -0.30 | -0.65 | 0.05 | -0.32 | -0.73 | 0.08 |
| mod_A * condition switch * session(^2) | -0.07 | -0.45 | 0.31 | -0.07 | -0.44 | 0.29 | -0.07 | -0.44 | 0.28 | -0.06 | -0.41 | 0.31 | -0.05 | -0.47 | 0.37 |
| condition single * mod_ch * session(^2) | -0.03 | -0.72 | 0.66 | -0.04 | -0.69 | 0.60 | -0.05 | -0.67 | 0.57 | -0.06 | -0.67 | 0.56 | -0.05 | -0.68 | 0.58 |
| condition switch * mod_ch * session(^2) | 0.08 | -0.62 | 0.78 | 0.07 | -0.60 | 0.73 | 0.07 | -0.58 | 0.70 | 0.10 | -0.51 | 0.73 | 0.13 | -0.54 | 0.79 |
